# Large-scale genomic characterisation of phage-plasmids in clinical Enterobacteriaceae isolates

**DOI:** 10.1101/2025.09.12.675607

**Authors:** Satheesh Nair, Clare R Barker, Matthew Bird, Alice Ledda, Caitlin Collins, Ryan Morrison, David R Greig, Ella V Rodwell, Anaïs Painset, Adam Crewdson, Claire Jenkins, Marie A Chattaway, Xavier Didelot, Paolo Ribeca

## Abstract

The life cycle of certain bacteriophages involves their maintenance within the bacterial cell as extrachromosomal elements, complete with replication and partitioning systems. These “phage-plasmids” (P-Ps) are distributed widely among bacterial phyla but are not typically included during genomic surveillance studies, and previous reports do not consider the context of their host strain diversity.

We recently identified a P1-like P-P carrying the *bla*_CTX-M-15_ resistance gene in *Salmonella enterica* serovar Typhi, prompting a subsequent investigation into the overall frequency of P-Ps among gastrointestinal bacteria under routine genomic surveillance within England. We expanded our study to include P-P groups known to be associated with Enterobacteriaceae (P1, D6, SSU5, N15) and scanned a collection of 66,856 genomes of diarrhoeagenic *Escherichia* spp., *Shigella* spp. and *S. enterica*.

All four P-P groups were detected in our dataset, totalling 9% of *E. coli* and *Shigella* genomes and 2% of *S. enterica* genomes. A small subset harboured two distinct P-P groups. P1-like P-Ps, some of which carried a cytotoxic necrotising factor, were predominantly associated with Shiga toxin-producing *E. coli* strains of public health concern, including clonal complexes CC11 (O157:H7), CC29 (O26:H11) and CC165. In contrast, SSU5 group P-Ps were linked with multidrug-resistant lineages of both *S.* Typhi and *Shigella sonnei*.

We found multiple antimicrobial resistance genes inserted into P-Ps via transposons, integrons and insertion sequences, and numerous defence/anti-defence systems. There was evidence of vertical transmission but also many links across species, time and geography, indicating that horizontal transfer is occurring regularly.

Phage-plasmids are often described only as cryptic elements or not detected during genomic surveillance. We show that P-Ps are associated with clinically relevant lineages of human pathogens and can acquire accessory genes that may impact on disease severity and therefore should play a more prominent role in pathogen surveillance and epidemiology.

## Introduction

The adaptation and evolution of bacteria to novel challenges is facilitated by their ability to acquire genes through vehicles of horizontal gene transfer (HGT) such as plasmids, transposons and bacteriophages. Bacteriophages (phages) are not only pervasive in nature but are the planet’s most abundant biological entity and have co-evolved with bacteria along the aeons (1). Phages can either kill bacteria directly through a lytic process or persist and integrate into the bacterial chromosome to form a lysogen (temperate phages). Like plasmids, they are a driving force in HGT as they harbour genes responsible for toxins, virulence factors and even antimicrobial resistance (AMR) (1,2).

In some cases, temperate phages belonging to certain families harbour plasmid-like replication and partitioning genes allowing them to behave as low-copy-number, self-replicating extrachromosomal elements termed phage-plasmids (P-Ps). These elements have seen a recent increase in research attention due to their links with bacterial evolution (3,4). P-Ps are often misclassified as plasmids and, compared to phages and plasmids, P-Ps are systematically understudied (5,6), although they are found throughout diverse aquatic and terrestrial environments and are thought to shape the microbial niches in these environments (7): indeed, even the abundant crAssphage was recently determined to be a linear P-P (8). There is a growing interest in P-Ps as they are involved in the transmission of AMR, virulence, defence systems and other accessory traits such as auxiliary metabolic pathways (7,9–12).

Recent comprehensive studies of P-P diversity (5,6) have identified four main groups that are associated with the Enterobacteriaceae family of bacteria: the well-characterised, circular P1 and its relative (or subgroup) D6 (13–15); the linear ‘telomere phage’ N15 (16–18), which is related to the lambdoid phages (19,20); and those belonging to the SSU5 supergroup/supercluster (5,15), which comprises at least four groups and includes SSU5 itself (15,21,22), pHCM2 of *S.* Typhi (23) and the virulence ‘plasmid’ pMT1 of *Y. pestis* (24).

AMR genes encoding resistance to many classes of antibiotics – importantly including extended-spectrum β-lactamases (ESBLs) and colistin resistance genes – have been detected in numerous P-Ps from both the P1 and SSU5 groups (9,25–31). The majority of these P-Ps carrying AMR genes have been found in Enterobacteriaceae: in a recent large-scale study (9), most were found in the genomes of just four hosts: *Acinetobacter baumannii*, *Escherichia coli*, *Klebsiella pneumoniae*, and *Salmonella* (spp. and *enterica*), thereby indicating the importance of P-Ps within the Enterobacteriaceae and especially in clinically relevant species.

Our previous screening for P1-like P-Ps in a collection of ∼47,000 *S. enterica* genomes deriving from human disease (11,31) showed that P-Ps belonging to the P1 group were more widely present in Salmonellae than previously thought, which highlighted the importance of gaining a better knowledge of P-P prevalence in broader datasets of clinically significant bacterial pathogens. Therefore, we set out in the current work to generalise and expand our analysis to include the four P-P groups (P1, D6, SSU5 and N15) within a larger collection of clinical Enterobacteriaceae isolates. This dataset additionally covers ∼19,000 *Shigella* and *Escherichia* spp. – predominantly the Shiga toxin-producing *E. coli* pathovar (STEC) but also other pathovars and species responsible for gastrointestinal infections – genomes that have been received and sequenced by the UK national reference laboratory for gastrointestinal bacteria.

We aimed to: analyse the distribution of the four P-P groups within the population structure of these pathogens; examine the genetic diversity of the P-P sequences, with a particular view to accessory genes; explore the relationship between horizontal and vertical gene transfer, and assess how spillover across different Enterobacteriaceae species due to HGT might engender the transmission of potentially virulent genes; and characterise the distribution of defence/anti-defence genes. For clarity we have reported on each of the P-P groups separately, as there was extensive variability among the different groups.

## Results

Our initial literature search for P-P groups likely to be present in the gastrointestinal pathogens included in our dataset (namely *Escherichia, Shigella* and *Salmonella* species) led to a focus on the P1, D6, SSU5 and N15 groups. To characterise the genetic variability across these groups and determine genes of interest we downloaded and investigated the reference sequences for these P-Ps (Fig. 1). We noted that, despite their shared nature as extrachromosomal elements, the P-P groups had distinct genomic structures that were well conserved within, but not between, the groups (Fig. 1A), and were only distantly related when highlighted on a prokaryotic viral taxonomy (Fig. 1B). From this analysis, we selected marker genes to represent each of the groups (highlighted in Fig. 1A), and proceeded to screen a total of 66,856 *Escherichia*, *Shigella* and *Salmonella* spp. isolate genomes for these marker genes.

**Figure 1.**
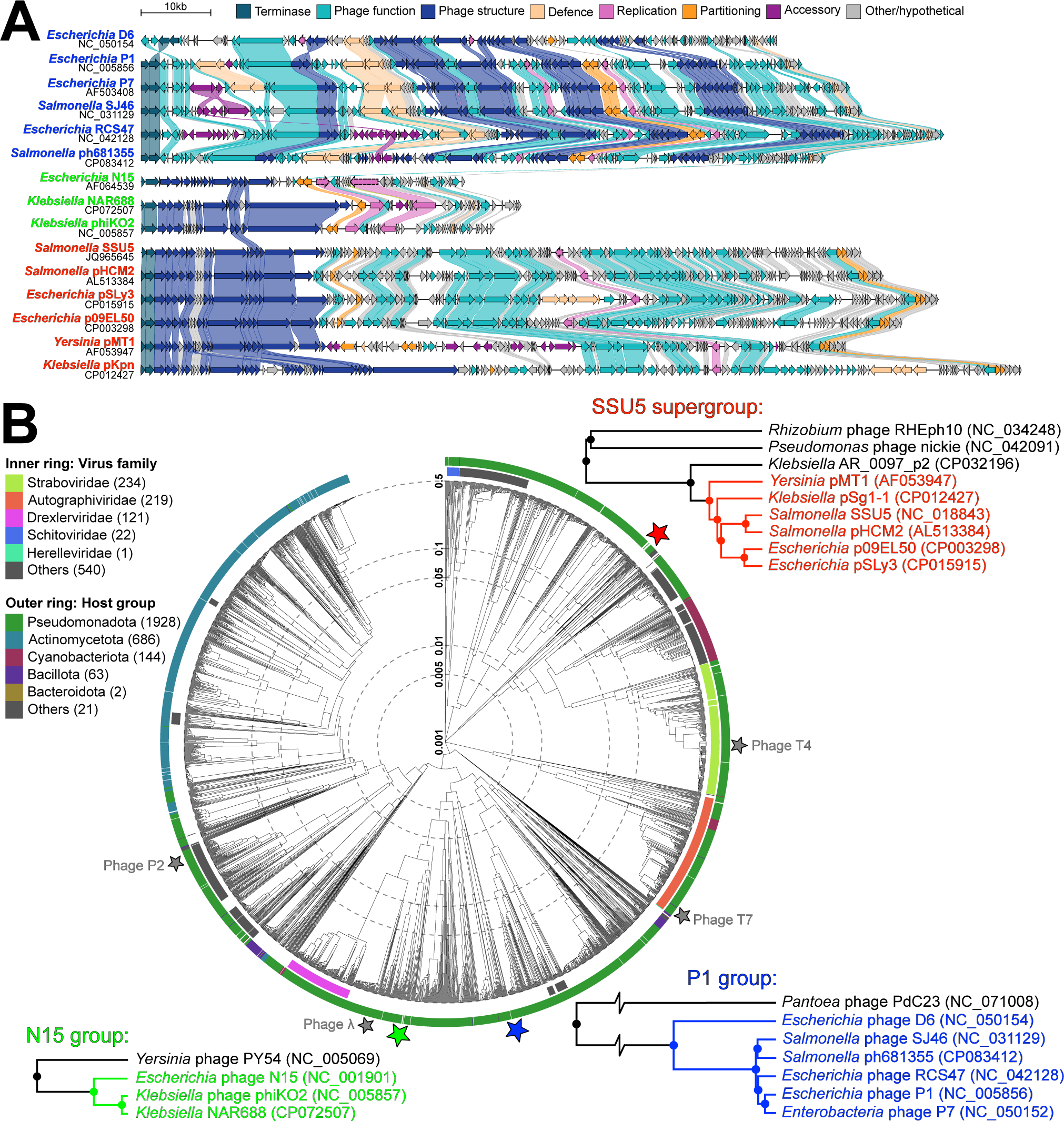
Taxonomic context and genetic diversity of the three phage-plasmid families present in Enterobacteriaceae. **A)** Comparison of reference sequences from the P-P groups considered in this work (P1, D6, N15, and SSU5) and members of their subgroups, showing similarities and differences in size, layout and gene content, with genes coloured by function and linked where protein sequence identity is >30%. Genes outlined in bold dashed lines are those chosen for screening. **B)** Dendrogram showing the locations of the P-Ps in relation to other related bacterial dsDNA viruses, based on tBLASTx genome-wide sequence similarity. Surrounding are more detailed views of the immediate relatives of the reference genome for each group, with selected P-Ps from previous studies.

Overall, we detected 2,641 P-Ps across 2,584 (3.9%) of the genomes, with frequency dependent on the host; ranging from 1.7% of *Salmonella* isolates to 8.4% in *Escherichia* and 10.4% in *Shigella* (Table 1, Fig. 2). Some of the most common organisms within our dataset had notably low P-P occurrence: clonal complex (CC) 11 (STEC O157:H7), accounting for a third of all *E. coli* isolate genomes, had only 141 positives and so represented <15% of the P-Ps in *E. coli*; *S. enterica* serovar Enteritidis corresponds to 27% of the *S. enterica* dataset, but only eight isolates (0.06%) had P-P sequences; and *Shigella flexneri*, despite making up 45% of the *Shigella* collection, had a prevalence of 33 positive isolates (or 4.4% of all P-Ps in *Shigella*). In addition, several of the largest groups within *E. coli* (including ST442 and ST738) and *S. enterica* (such as *S.* Chester and *S.* Bovismorbificans) were entirely lacking P-Ps. Meanwhile, certain species (*Shigella sonnei*, *Escherichia albertii*), *S. enterica* serovars (*S.* Typhi), and *E. coli* ST/CCs (CC29, CC165, CC28, CC131 and ST342) had notably high prevalence (Fig. 2A). Among the positive isolates, 43 *Escherichia* spp., nine *S. sonnei* and five *S. enterica* genomes possessed two different types of P-P, with this phenomenon being particularly common in *E. coli* CC165: 22% of CC165 isolates with P-Ps had two groups present.

**Figure 2.**
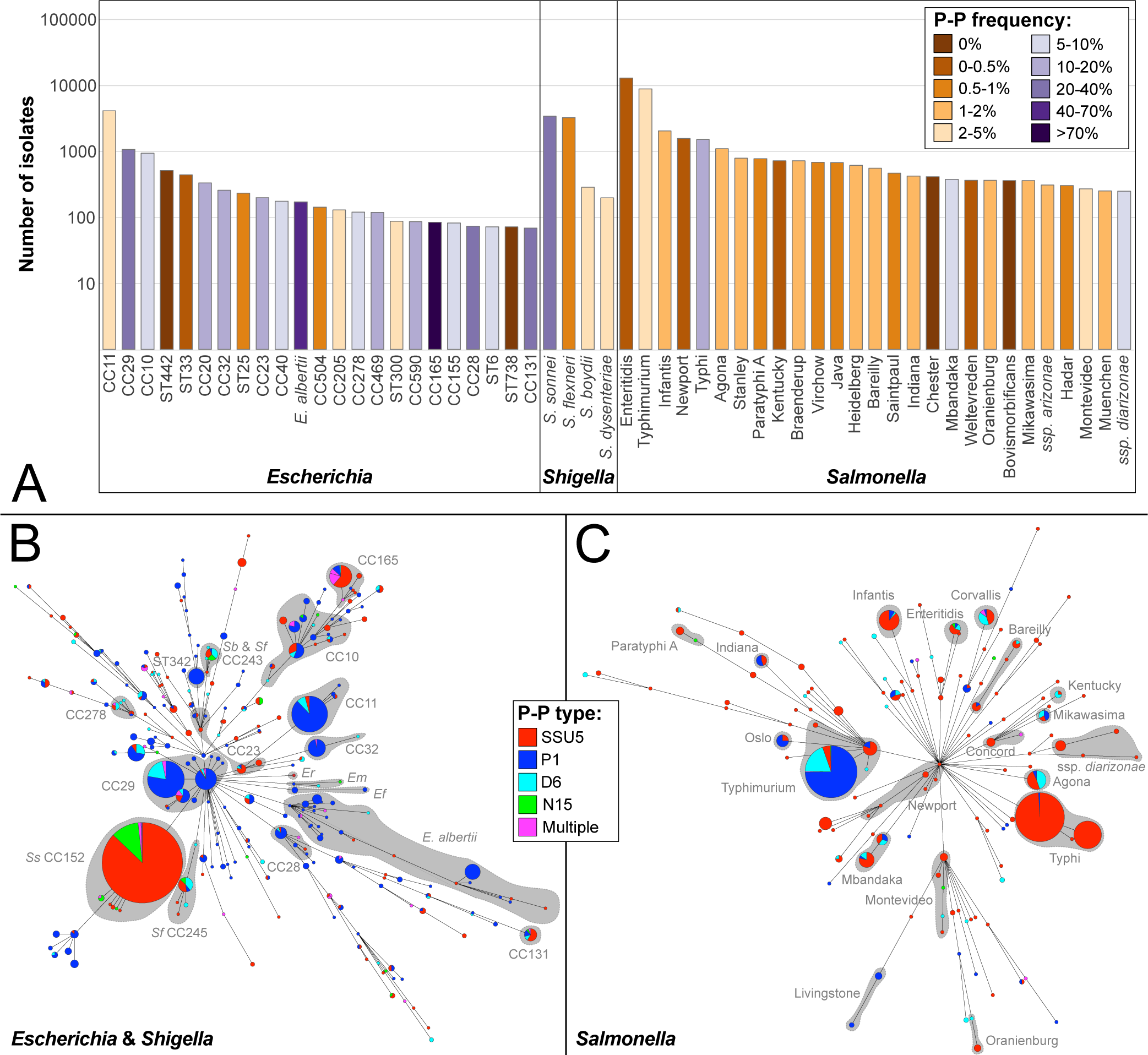
Phage-plasmid prevalence among UK *Escherichia*, *Shigella* and *Salmonella* isolates. **A)** Number of isolates belonging to the most common organisms within the overall dataset, coloured by frequency of P-P carriage within that group (ST, sequence type; CC, clonal complex). **B)** Minimum-spanning tree produced from MLST profiles of the *Escherichia* spp. & *Shigella* spp. genomes containing P-Ps, coloured by P-P group with main ST/CC or species labelled (Ef, *E. fergusonii*; Em, *E. marmotae;* Er, *E. ruysiae*; Sb, *S. boydii*; *Sf*, *S. flexneri*). **C)** Minimum-spanning tree produced from MLST profiles of *S. enterica* genomes containing P-Ps, coloured by P-P group and labelled with main serovars or subspecies.

**Table 1.** Distribution of P-Ps in isolates from species and subspecies of *Escherichia*, *Shigella* and *Salmonella*, including serovars and sequence types (STs)/clonal complexes (CCs) containing >20 positive isolates. Frequency for *E. fergusonii*, *E. marmotae* and *E. ruysiae* not included due to low sample sizes. Expanded version available as Supplementary Table S3.

| Group | P1 | D6 | SSU5 | N15 | P1+<br>D6 | P1+<br>SSU5 | SSU5<br>+D6 | SSU5<br>+N15 | Total<br>no. | Overall<br>freq. |
| --- | --- | --- | --- | --- | --- | --- | --- | --- | --- | --- |
| ***Escherichia* total** | **666** | **95** | **209** | **12** | **11** | **22** | **10** | **0** | **1025** | **8.4%** |
| *E. albertii* | 61 | 1 | 6 | 1 |  | 2 |  |  | 71 | 41.3% |
| *E. fergusonii* | 4 |  |  |  |  |  |  |  | 4 | - |
| *E. marmotae* |  | 1 |  | 1 |  |  |  |  | 2 | - |
| *E. ruysiae* |  |  |  |  |  |  | 1 |  | 1 | - |
| *E. coli* CC10 | 37 | 5 | 25 | 2 |  | 3 |  |  | 72 | 7.7% |
| *E. coli* CC11 | 126 | 9 | 6 |  |  |  | 1 |  | 141 | 3.4% |
| *E. coli* CC20 | 23 | 9 | 4 | 1 |  | 1 |  |  | 38 | 11.4% |
| *E. coli* CC23 | 4 | 2 | 15 |  |  |  |  |  | 21 | 10.5% |
| *E. coli* CC29 | 183 | 33 | 6 |  | 8 | 1 |  |  | 231 | 21.6% |
| *E. coli* CC32 | 31 |  |  |  |  |  |  |  | 31 | 12.1% |
| *E. coli* CC165 | 7 | 1 | 38 |  |  | 9 | 4 |  | 59 | 70.2% |
| *E. coli* ST342 | 27 |  |  |  |  |  |  |  | 27 | 57.4% |
| Other ST/CC |  |  |  |  |  |  |  |  | 326 | - |
| ***Shigella* total** | **5** | **16** | **630** | **88** | **0** | **0** | **0** | **9** | **748** | **10.4%** |
| *S. sonnei* |  |  | 616 | 81 |  |  |  | 9 | 706 | 20.6% |
| *S. flexneri* | 1 | 13 | 11 | 7 |  |  |  |  | 32 | 1.0% |
| *S. dysenteriae* | 3 | 1 |  |  |  |  |  |  | 4 | 2.1% |
| *S. boydii* | 1 | 2 | 3 |  |  |  |  |  | 6 | 2.0% |
| ***Salmonella* total** | **216** | **96** | **492** | **2** | **0** | **1** | **4** | **0** | **811** | **1.7%** |
| *S. Infantis* | 3 | 1 | 27 |  |  |  |  |  | 31 | 1.5% |
| *S. Mbandaka* | 4 | 6 | 19 |  |  |  |  |  | 29 | 7.7% |
| *S. Typhi* | 2 |  | 254 |  |  |  |  |  | 256 | 16.9% |
| *S. Typhimurium* | 166 | 43 | 22 | 1 |  | 1 |  |  | 233 | 2.6% |
| Other serovar | 41 | 46 | 147 | 1 |  |  | 4 |  | 239 | - |
| ssp. arizonae |  |  | 6 |  |  |  |  |  | 6 | 1.9% |
| ssp. diarizonae |  |  | 14 |  |  |  |  |  | 14 | 5.6% |
| ssp. houtenae |  |  | 1 |  |  |  |  |  | 1 | 1.7% |
| ssp. salamae |  |  | 2 |  |  |  |  |  | 2 | 2.7% |
| **TOTAL** | **887** | **207** | **1331** | **102** | **11** | **23** | **14** | **9** | **2584** | **3.9%** |

### P1/D6 Group

We detected P1 and D6 group P-Ps in 921 and 232 isolates in total, respectively, with eleven *E. coli* isolates carrying both types. Within our *Escherichia* collection the P1-like P-Ps were most frequent among isolates belonging to CC165 (70.2% of genomes), ST342 (57.4%), *E. albertii* (41.3%), CC28 (25.7%) and CC29 (21.6%). P1-like P-Ps were rarely found in the *Shigella* spp. genomes: we observed five positive isolates from three species, at a total frequency of 0.07%.

To determine the distribution of P1-like P-Ps within the largest and most clinically relevant types of diarrhoeagenic *Escherichia* spp., and to investigate any association with host population structure, we produced phylogenetic trees for the two main STEC lineages (CC11 O157:H7 and CC29 O26:H11) and the five other groups within *Escherichia* with >20% P-P frequency (*E. albertii*, CC165, CC131, CC28 and ST342), and plotted P-P distribution among these hosts (Fig.S1-S4). Together, these seven host groups accounted for 49% of the P1-like P-P positive isolates, or 64% of those in the *Escherichia* dataset. The P1-like P-Ps were distributed sporadically across the whole of the CC29 population, however in CC11, CC165 and *E. albertii* they appeared more frequently in particular clades; being most associated with sub-lineages Ic and IIa of CC11, with ST301 of CC165, and with avian clusters 6 and 7 of *E. albertii*. Within CC165, all but one of the P1-like P-Ps belonged to ST301 isolates, which is a sub-lineage with high frequency of AMR and virulence genes(32,33): in fact, 95% of the ST301 isolates in this dataset carried at least one P-P element.

As we described previously(11), in *S. enterica* the P1-like P-Ps were present at a low frequency (<0.5%) across more than 20 different serovars and were most common in *S.* Typhimurium eBG1 (n=167, 1.9%), *S.* Oslo (n=9, 6.1%) and *S.* Livingstone (n=6, 6.1%). Our results here show that the D6-like P-Ps were also rare in *S. enterica* (n=100, 0.21%) but were again most numerous in *S.* Typhimurium eBG1 (n=43, 0.5%); particularly ST34 (associated with the monophasic variant of *S.* Typhimurium) but also including one *S.* Typhimurium belonging to the emergent invasive, multidrug-resistant (MDR) ST213 group(34). The D6-like P-Ps were most frequent in *S.* Corvallis (n=9, 4.5%), *S.* Mbandaka (n=6, 1.6%) and *S.* Agona (n=12, 1.1%), and were otherwise distributed more sporadically across 19 other serovars. In *Escherichia* and *Shigella* spp. the D6-like P-Ps were also present widely at a low frequency and were most numerous in the largest STEC complexes – CC11 (n=9, 0.2%) and CC29 (n=41, 3.4%) – as well as in CC20 (n=9, 2.7%) and *S. flexneri* (n=13, 0.4%). Interestingly, there were also singular isolates of *E. albertii* and *E. marmotae* with a D6-like P-P, as well as a small number of *S. boydii* and *S. dysenteriae*.

To examine the diversity and population structure of the P1/D6 group P-Ps present in our dataset, we generated a tree containing the extracted P-P contigs alongside related publicly available element genomes (Fig. 3). Within this, several clades were comprised largely of P-Ps from a single host group (such as CC11, CC29, *E. albertii* or *S.* Typhimurium eBG1), suggesting that these elements are being maintained by vertical inheritance in certain populations. However, the overall tree did not match the population structure of the Enterobacteriaceae, indicative of frequent horizontal transfer of these elements. In support of this, within the P1/D6 group tree we observed several instances of identical or closely related P-P sequences that span multiple CCs/serovars or species, as well as time and geography (Fig. S5).

**Figure 3.**
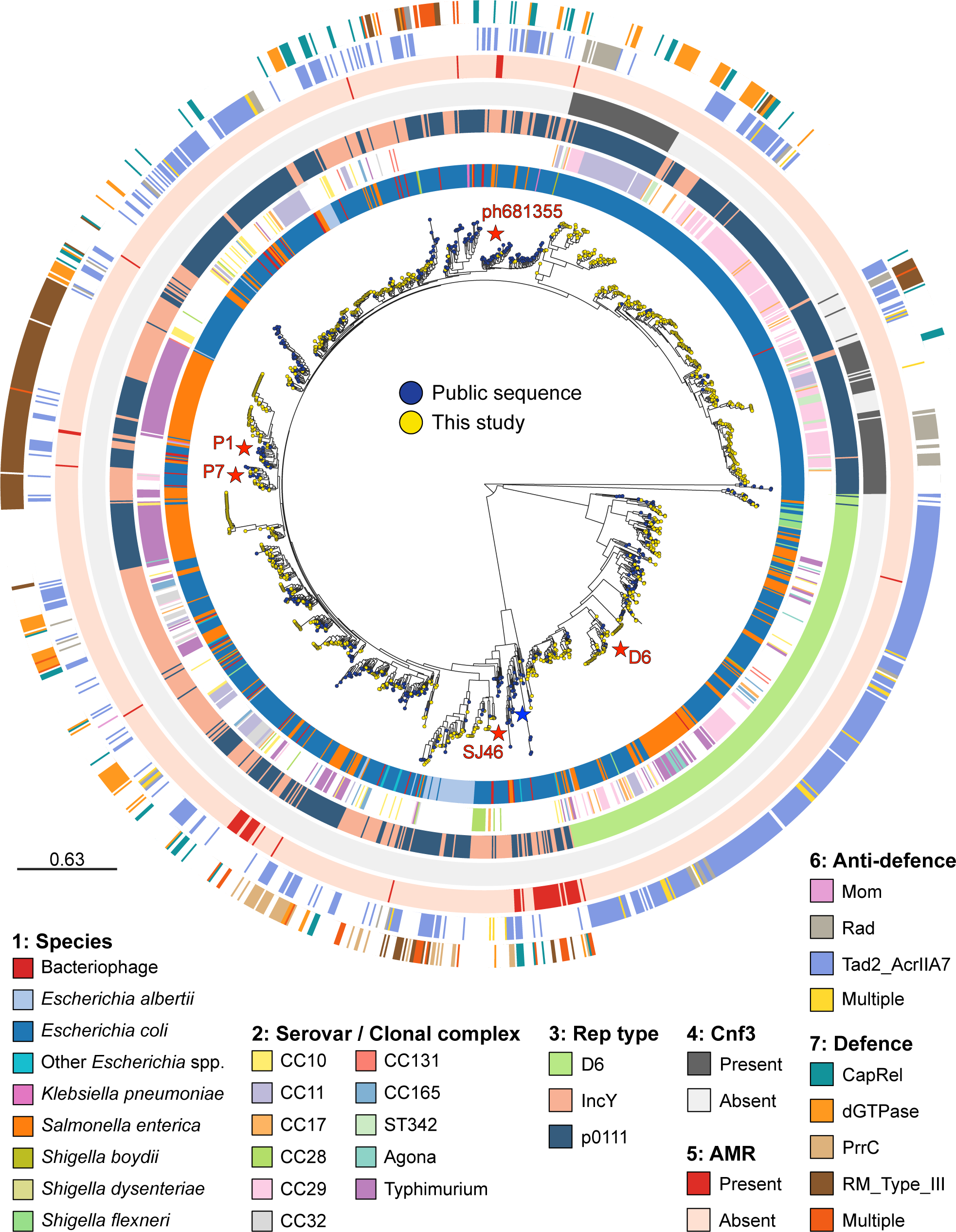
Midpoint-rooted tree showing the population structure of the P1/D6 group including public sequences (n=1592 total), with metadata of bacterial host strain where available. Main serovars (*Salmonella*) and clonal complexes (*Escherichia*) are shown, the location of references and sequences of interest are highlighted in red, and the AMR clade is highlighted with a blue star. Defence and anti-defence systems are shown with the exception of the Dar-Ddr system, as it was found to be universally present in P1 and D6 P-Ps (Table 3).

To examine accessory genes of relevance to public health within the P1 and D6 group elements present in our collection, we screened the contigs for the presence of AMR genes. Alongside the two *S.* Typhi P1-like elements carrying IS*Ec9*-encoded (synonym IS*Ecp1*) *bla*_CTX-M-15_ on which we previously reported (11,31), we also detected AMR genes on P-Ps in five other isolates (Table 2). Four of these were also P1-like P-Ps, all deriving from *E. coli* (CC10 and CC20 only), which carried various genes including *tet(A)*, *bla*_TEM-1_ and *bla*_CTX-M-15_. The singular D6-like P-P with AMR belonged to an *S.* Kentucky isolate and showed a partial, end-of-contig match to the *bla*_TEM-1_ gene (100% identity, 51.4% coverage). We also noted from the tree that AMR genes present in public sequences appeared to be more frequent in certain clades (Fig. 3). In particular, the clade lying basal to the main P1 lineage that contained AMR-carrying P-Ps from two UK CC10 isolates, alongside public P-Ps deriving from globally distributed *E. coli* belonging to CC165, CC23 and several other STs (Fig. S6).

**Table 2.** AMR genes detected in D6, P1 and SSU5 P-Ps, and associated metadata including where host isolates belong to a 10-SNP cluster (Table S2). * = described previously as an AMR plasmid in patients with travellers’ diarrhoea (37). ª = shown in Figure 5A. ^b^ = shown in Figure 5B. ^c^ = shown in Figure 5C. ^d^ = shown in Figure 5D. ^x^ = no contig available. ^1^ = in AMR clade 1. ^2^ = in AMR clade 2.

| Isolate | Strain | CC/eBG | Cluster | Year | P-P | Gene(s) | Element | Site |
| --- | --- | --- | --- | --- | --- | --- | --- | --- |
| 737225 | S. Kentucky | eBG56 |  | 2019 | D6 | *bla*<sub>TEM-1</sub> | Unknown | - |
| 206715 | *E. coli* O89:H5 | CC20 |  | 2016 | P1 | *bla*<sub>CTX-M-15</sub> | ISEc9 | - |
| 548299 | *E. coli* O25:H40 | CC10 |  | 2018 |  | *aadA1; dfrA1; merCRTP; qacE; sul1; tet(A)* | Tn3; IS1236 | - |
| 1493978 | *E. coli* O123:H40 | CC10 |  | 2021 |  | *bla*<sub>TEM-1</sub>; *tet(A)* | Tn3 | - |
| 637692 | *E. coli* O166:H25 | CC10 |  | 2018 |  | *merRTP* | Unknown | - |
| 681355 | S. Typhi | eBG13 | Typhi1 | 2019 |  | *bla*<sub>CTX-M-15</sub> | ISEc9 | - |
| 672572 | S. Typhi | eBG13 | Typhi1 | 2019 |  | *bla*<sub>CTX-M-15</sub> | ISEc9 | - |
| 660450<sup>bx</sup> | *E. albertii* | ST4634 | EA1 | 2018 | SSU5-pSLy3 | *bla*<sub>TEM-135</sub> | Tn3 | 3 |
| 313858<sup>bx</sup> | *E. coli* O21:H21 | CC40 |  | 2016 |  | *bla*<sub>TEM-1</sub> | Tn3 | 4 |
| 868390<sup>c2</sup> | *E. coli* O33:H16 | ST295 |  | 2019 |  | *bla*<sub>CTX-M-15</sub> | ISEc9 | 2 |
| 899047<sup>x</sup> | *E. coli* O8:H9 | CC23 | CC23a | 2019 |  | *bla*<sub>CTX-M-15</sub> | ISEc9 | 2 |
| 794726<sup>x</sup> | *E. coli* O8:H9 | CC23 |  | 2019 |  | *bla*<sub>CTX-M-15</sub> | ISEc9 | 2 |
| 255516<sup>ac2</sup> | *E. coli* O25:H42 | CC10 |  | 2016 |  | *bla*<sub>CTX-M-15</sub> | ISEc9 | 2 |
| 296627<sup>x</sup> | *E. coli* O131:H25 | CC40 |  | 2016 |  | *bla*<sub>CTX-M-15</sub> | ISEc9 | 2 |
| 822583<sup>c2</sup> | *E. coli* O33:H34 | ST2346 |  | 2019 |  | *bla*<sub>CTX-M-15</sub> | ISEc9 | 2 |
| 877250<sup>c2</sup> | *E. coli* O44:H18 | ST394 |  | 2020 |  | *bla*<sub>CTX-M-15</sub> | ISEc9 | 8 |
| 256169<sup>2</sup> | *E. coli* O85:H9 | CC590 |  | 2016 |  | *bla*<sub>CTX-M-15</sub> | ISEc9 | 8 |
| 654720<sup>x</sup> | *E. coli* O170:H2 | CC165 |  | 2018 |  | *bla*<sub>CTX-M-15</sub> | ISEc9 | 8 |
| 773772<sup>2</sup> | *E. coli* O?:H5 | CC165 | CC165d | 2019 |  | *bla*<sub>CTX-M-15</sub> | ISEc9 | 8 |
| 1078073<sup>2</sup> | *E. coli* O?:H5 | CC165 |  | 2021 |  | *bla*<sub>CTX-M-15</sub> | ISEc9 | 8 |
| 897289<sup>c2</sup> | *E. coli* O55:H21 | CC4213 |  | 2020 |  | *bla*<sub>CTX-M-15</sub> | ISEc9 | 8 |
| 637747<sup>2</sup> | *E. coli* O89:H5 | CC20 |  | 2018 |  | *bla*<sub>CTX-M-15</sub> | ISEc9 | 8 |
| 723831<sup>2</sup> | *S. flexneri* 6 | CC145 |  | 2019 |  | *bla*<sub>CTX-M-15</sub> | ISEc9 | 8 |
| 435526<sup>2</sup> | *S. boydii* 2 | CC145 |  | 2017 |  | *bla*<sub>CTX-M-15</sub> | ISEc9 | 8 |
| 410905<sup>2</sup> | *S. flexneri* 6 | CC145 | Flex1 | 2017 |  | *bla*<sub>CTX-M-15</sub> | ISEc9 | 8 |
| 399473<sup>2</sup> | *S. flexneri* 6 | CC145 | Flex1 | 2017 |  | *bla*<sub>CTX-M-15</sub> | ISEc9 | 8 |
| 797720<sup>c2</sup> | *E. coli* O25:H16 | ST1491 |  | 2019 |  | *bla*<sub>CTX-M-15</sub> | ISEc9 | 1 |
| 388744<sup>c2</sup> | *E. coli* O25:H16 | ST1491 |  | 2017 |  | *bla*<sub>CTX-M-15</sub> | ISEc9 | 1 |
| 363476<sup>2</sup> | *E. coli* O111:H19 | CC469 |  | 2017 |  | *bla*<sub>CTX-M-15</sub> | ISEc9 | 1 |
| 912518<sup>2</sup> | *E. coli* O89:H45 | CC10 |  | 2020 |  | *bla*<sub>CTX-M-15</sub> | ISEc9 | 1 |
| 533373<sup>c2</sup> | *E. coli* O6:H10 | CC10 |  | 2018 |  | *bla*<sub>CTX-M-15</sub> | ISEc9 | 1 |
| 288997<sup>c2</sup> | *E. coli* O25:H17 | CC131 |  | 2016 |  | *bla*<sub>CTX-M-15</sub> | ISEc9 | 1 |
| 894913<sup>c2</sup> | *S. flexneri* 2a | CC245 | Flex5 | 2020 |  | *bla*<sub>CTX-M-15</sub> | ISEc9 | 1 |
| 951667<sup>c2</sup> | *S. flexneri* 2a | CC245 | Flex5 | 2020 |  | *bla*<sub>CTX-M-15</sub> | ISEc9 | 1 |
| 946587<sup>c2</sup> | *S. flexneri* 2a | CC245 | Flex5 | 2020 |  | *bla*<sub>CTX-M-15</sub> | ISEc9 | 1 |
| 946622<sup>ac2</sup> | *S. flexneri* 2a | CC245 | Flex5 | 2020 |  | *bla*<sub>CTX-M-15</sub> | ISEc9 | 1 |
| 627377<sup>c2</sup> | *S. flexneri* 2a | CC245 |  | 2018 |  | *bla*<sub>CTX-M-15</sub> | ISEc9 | 1 |
| 788309<sup>*ac</sup> | *E. coli* O?:H31 | ST3032 |  | 2019 |  | *bla*<sub>CTX-M-15</sub> | ISEc9 | 5 |
| 246161<sup>c2</sup> | *E. coli* O6:H10 | CC10 | CC10b | 2016 |  | *bla*<sub>CTX-M-15</sub> | ISEc9 | 6 |
| 248618<sup>c2</sup> | *E. coli* O6:H10 | CC10 | CC10b | 2016 |  | *bla*<sub>CTX-M-15</sub> | ISEc9 | 6 |
| 738601<sup>ac</sup> | *E. coli* O6:H10 | CC10 |  | 2019 |  | *bla*<sub>CTX-M-15</sub> | ISEc9 | 6 |
| 872085<sup>2</sup> | *S. boydii* 1 | CC145 |  | 2020 |  | *bla*<sub>CTX-M-15</sub> | ISEc9 | 7 |
| 524337<sup>c2</sup> | *E. coli* O?:H19 | CC469 |  | 2018 |  | *bla*<sub>CTX-M-15</sub> | ISEc9 | 7 |
| 738636<sup>2</sup> | *E. coli* O167:H5 | CC205 | CC205a | 2019 |  | *bla*<sub>CTX-M-15</sub> | ISEc9 | 7 |
| 899037<sup>*ac2</sup> | *E. coli* O167:H5 | CC205 | CC205a | 2019 |  | *bla*<sub>CTX-M-15</sub> | ISEc9 | 7 |
| 358046<sup>2</sup> | *S. sonnei* | CC152 |  | 2017 |  | *bla*<sub>CTX-M-15</sub> | ISEc9 | 7 |
| 950327 | *S. Typhimurium* | eBG1 | Typhim5 | 2020 | SSU5-pHCM2 | *bla*<sub>TEM-1</sub> | Tn3 | 4 |
| 267046<sup>a</sup> | *S. Worthington* | eBG101 |  | 2016 |  | *merADERT* | Unknown | 3 |
| 804434<sup>a1</sup> | *S. Typhimurium* | eBG1 | Typhim7 | 2019 |  | *aadA3; dfrA21; qacE; sul1* | Int1/IS6100 | 4 |
| 790484<sup>1</sup> | *S. Typhimurium* | eBG1 | Typhim7 | 2019 |  | *aadA3; dfrA21; qacE; sul1* | Int1/IS6100 | 4 |
| 816218<sup>1</sup> | *S. Montevideo* | eBG40 |  | 2019 |  | *dfrA21; qacE; sul1* | Int1/IS6100/Tn3 | 4 |
| 235032<sup>1</sup> | *S. Mbandaka* | eBG62 |  | 2016 |  | *dfrA15; qacE; sul1* | Int1/IS6100/Tn3 | 4 |

Within our collection were seven STEC isolates for which we had complete, closed genomes available from long-read sequencing projects. A manual examination of the circular P1 sequences from these genomes showed that four (two each of CC11 and CC29) carried a gene annotated as *cnf3* (cytotoxic necrotising factor)(35,36) adjacent to insertion sequence (IS*66*) transposase genes. We then scanned the complete P1 contig collection, including the downloaded public sequences, for the presence of this *cnf3* gene (n=143 positive) and found that it was restricted to a single clade within the P1 P-Ps (Fig. 3). The *cnf3*-associated clade contained 338 genomes in total and primarily comprised of elements from *E. coli* CC29 O26:H11 (175/338, 52%), CC11 O157:H7 (65/338, 19%) and ST342 (21/338, 6%) isolates. Within this clade, the 143 P-Ps found to be positive for *cnf3* again mostly belonged to either CC11 (65/65), CC29 (48/175) or ST342 (16/21), and there were two public P1-like sequences with *cnf3* (CP045829 – human, Australia 2015; CP114131 – pig, China 2018) whose whole genome assemblies indicated that these also originated from CC11 and CC29 host strains. There were no AMR genes present in this clade, and 90.5% had the p0111 replication gene, rather than IncY. A search of sequences within GenBank, and those from our collection without P-Ps, showed that P1-like P-Ps carrying *cnf3* were spread globally and associated with multiple hosts, and that *cnf3* could also be found on conjugative plasmids or inserted into the chromosome, where interestingly it was located <50 kbp from a Shiga toxin-encoding prophage in at least one STEC genome (Fig. S7A). As the impact of the Cnf3 toxin on human pathogenicity is not yet well described, we compared its amino acid sequence to other Cnf variants, which have proven cytotoxic affects both *in vitro* and *in vivo* (36). Cnf3 shared 70% identity with Cnf1 and Cnf2 (both also deriving from *E. coli*), and 68% with CnfY of *Y. pseudotuberculosis* (Fig. S7B).

Alongside the virulence and resistance typing, we also screened the extracted P1/D6 group sequences for the presence of phage defence and anti-defence systems (Table 3, Table 4). The anti-RM (restriction-modification) ‘defence against restriction’ system (comprising DarAB, DdrAB, Hdf and Ulx), which is very well described in P1 (38,39), was present in every sequence – though *ddrB* was missing from all but four of the D6 sequences. We also detected the Tad2 AcrIIA7 (anti-Thoeris) system in the majority of sequences (93% D6, 56% P1). The Rad (anti-Retron) system (largely restricted to two clades), Abc2 (anti-RecBCD) and Mom (anti-RM) systems were also present, in lower numbers.

**Table 3.**
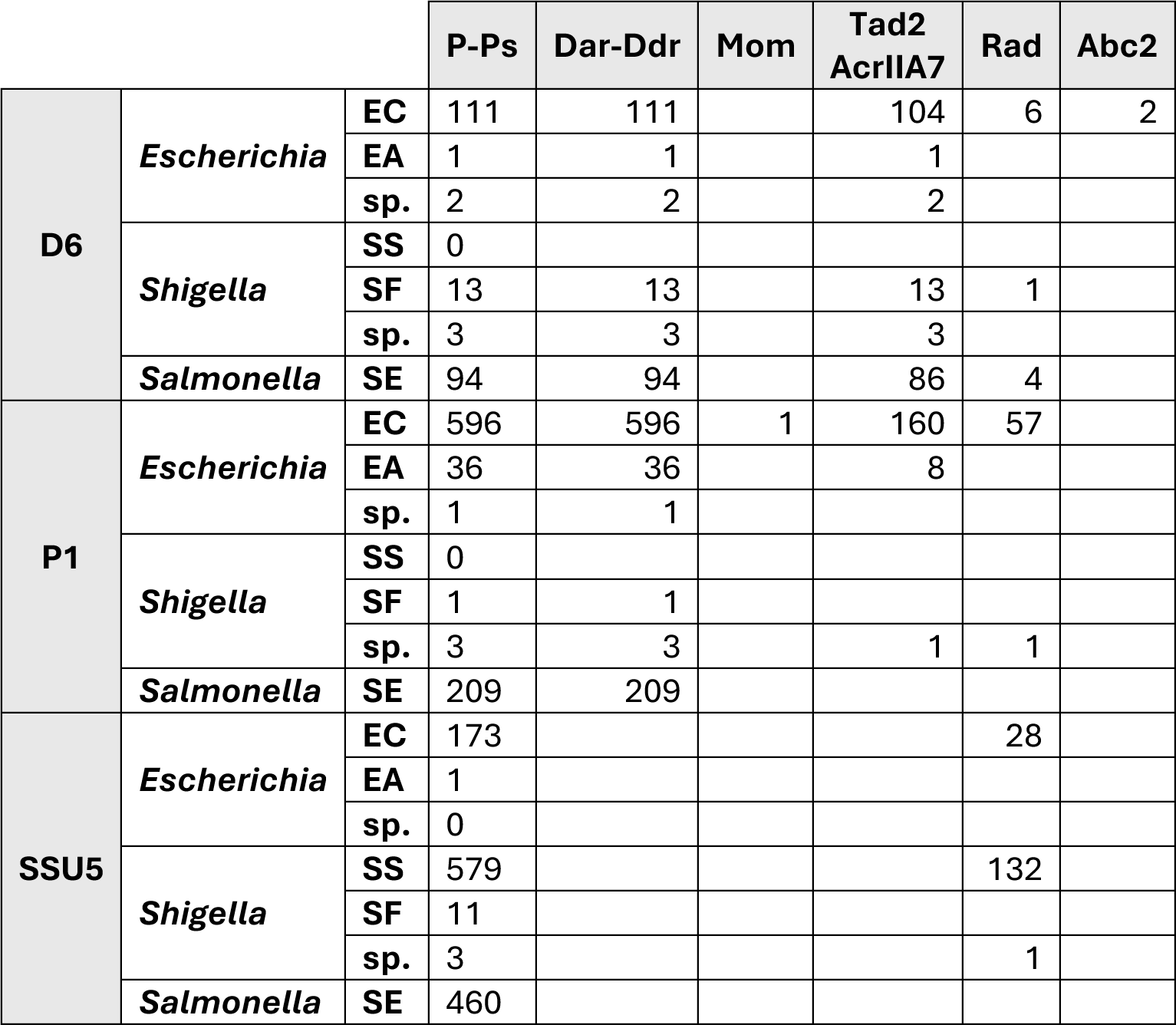
Summary of complete anti-defence systems found in P-P contigs, sorted by bacterial genera, species, and P-P group. Abbreviations: EA, *E. albertii*; EC, *E. coli*; SD, *S. dysenteriae*; SF, *S. flexneri*; SS, *S. sonnei*; SE, *S. enterica*.

|  |  |  | P-Ps | Dar-Ddr | Mom | Tad2<br>AcrIIA7 | Rad | Abc2 |
| --- | --- | --- | --- | --- | --- | --- | --- | --- |
| D6 | *Escherichia* | EC | 111 | 111 |  | 104 | 6 | 2 |
|  |  | EA | 1 | 1 |  | 1 |  |  |
|  |  | sp. | 2 | 2 |  | 2 |  |  |
|  | *Shigella* | SS | 0 |  |  |  |  |  |
|  |  | SF | 13 | 13 |  | 13 | 1 |  |
|  |  | sp. | 3 | 3 |  | 3 |  |  |
|  | *Salmonella* | SE | 94 | 94 |  | 86 | 4 |  |
| P1 | *Escherichia* | EC | 596 | 596 | 1 | 160 | 57 |  |
|  |  | EA | 36 | 36 |  | 8 |  |  |
|  |  | sp. | 1 | 1 |  |  |  |  |
|  | *Shigella* | SS | 0 |  |  |  |  |  |
|  |  | SF | 1 | 1 |  |  |  |  |
|  |  | sp. | 3 | 3 |  | 1 | 1 |  |
|  | *Salmonella* | SE | 209 | 209 |  |  |  |  |
| SSU5 | *Escherichia* | EC | 173 |  |  |  | 28 |  |
|  |  | EA | 1 |  |  |  |  |  |
|  |  | sp. | 0 |  |  |  |  |  |
|  | *Shigella* | SS | 579 |  |  |  | 132 |  |
|  |  | SF | 11 |  |  |  |  |  |
|  |  | sp. | 3 |  |  |  | 1 |  |
|  | *Salmonella* | SE | 460 |  |  |  |  |  |

**Table 4.** Summary of complete defence systems found in P-P contigs, sorted by bacterial genera, species, and P-P group. Abbreviations: EA, *E. albertii*; EC, *E. coli*; SD, *S. dysenteriae*; SF, *S. flexneri*; SS, *S. sonnei*; SE, *S. enterica*.

|  |  |  | P-Ps | PrrC | RM Type I | RM Type III | dGTPase | CapRel | PD-T4-3 |
| --- | --- | --- | --- | --- | --- | --- | --- | --- | --- |
| D6 | *Escherichia* | EC | 111 | 1 |  |  |  |  |  |
|  |  | EA | 1 |  |  |  |  |  |  |
|  |  | sp. | 2 |  |  |  |  |  |  |
|  | *Shigella* | SS | 0 |  |  |  |  |  |  |
|  |  | SF | 13 |  |  |  |  |  |  |
|  |  | sp. | 3 |  |  |  |  |  |  |
|  | *Salmonella* | SE | 94 |  |  |  |  |  |  |
| P1 | *Escherichia* | EC | 596 | 15 |  | 19 | 77 | 49 |  |
|  |  | EA | 36 | 9 |  | 12 |  |  |  |
|  |  | sp. | 1 |  |  |  |  |  |  |
|  | *Shigella* | SS | 0 |  |  |  |  |  |  |
|  |  | SF | 1 |  |  |  |  |  |  |
|  |  | sp. | 3 |  |  | 1 |  |  |  |
|  | *Salmonella* | SE | 209 | 17 |  | 87 | 5 | 3 |  |
| SSU5 | *Escherichia* | EC | 173 | 7 | 2 |  |  |  | 38 |
|  |  | EA | 1 |  |  |  |  |  | 1 |
|  |  | sp. | 0 |  |  |  |  |  |  |
|  | *Shigella* | SS | 579 |  |  |  |  |  | 13 |
|  |  | SF | 11 |  |  |  |  |  | 7 |
|  |  | sp. | 3 |  |  |  |  |  |  |
|  | *Salmonella* | SE | 460 | 29 |  | 2 |  |  |  |
|  |  |  | P-Ps | PD-T4-3 | PD-T7-5 | AbiH | DS-33 | DS-37 | HEC-06 |
| D6 | *Escherichia* | EC | 111 |  |  |  |  |  |  |
|  |  | EA | 1 |  |  |  |  |  |  |
|  |  | sp. | 2 |  |  |  |  |  |  |
|  | *Shigella* | SS | 0 |  |  |  |  |  |  |
|  |  | SF | 13 |  |  |  |  |  |  |
|  |  | sp. | 3 |  |  |  |  |  |  |
|  | *Salmonella* | SE | 94 |  |  |  |  |  |  |
| P1 | *Escherichia* | EC | 596 |  |  |  |  |  |  |
|  |  | EA | 36 |  |  |  |  |  |  |
|  |  | sp. | 1 |  |  |  |  |  |  |
|  | *Shigella* | SS | 0 |  |  |  |  |  |  |
|  |  | SF | 1 |  |  |  |  |  |  |
|  |  | sp. | 3 |  |  |  |  |  |  |
|  | *Salmonella* | SE | 209 |  |  |  |  |  |  |
| SSU5 | *Escherichia* | EC | 173 | 38 | 7 |  |  |  |  |
|  |  | EA | 1 | 1 |  |  |  |  |  |
|  |  | sp. | 0 |  |  |  |  |  |  |
|  | *Shigella* | SS | 579 | 13 |  |  |  |  |  |
|  |  | SF | 11 | 7 |  |  |  |  |  |
|  |  | sp. | 3 |  |  |  |  |  |  |
|  | *Salmonella* | SE | 460 |  |  | 30 | 4 | 7 | 2 |

Meanwhile, defence systems were mostly associated with P1 elements: only one D6-like P-P from an *E. coli* isolate had a complete defence system. In the P1-like sequences, 20 defence systems were found in the *E. albertii* contigs (56%), 165 in *E. coli* (28%) and 116 in *S. enterica* (55%). The defences found in P1 elements were mostly composed of abortive infection mechanisms, including Type III RM systems, dGTPase, PrrC (Type I RM) and CapRel (toxin-antitoxin, TA) systems. We noted differences between the species, with P1-like P-Ps deriving from *E. coli* more commonly carrying CapRel or dGTPase systems, while most of those from *S. enterica* had RM type III systems. When considering the wider predicted defences, we found numerous putative RM type II systems in both D6 and P1 sequences, and associations between PD-Lambda-5 in D6 P-Ps and Druantia II nuclease systems in P1 P-Ps (Fig. S8). Interestingly, P1-like P-Ps from *S. enterica* had the highest overall diversity of defence systems, despite their generally sporadic distribution.

### SSU5 Supergroup

Phage-plasmids within the SSU5 supergroup were detected in 1,377 isolate genomes in total, the majority of which were either *S. sonnei* (n=625, 45.4% of SSU5 positive genomes) or *S. enterica* serovar Typhi (n=254, 18.3%) (Table 1). Of the 880 SSU5 P-Ps found in *Escherichia* and *Shigella* spp., more than two thirds derived from *S. sonnei*: 18% of all *S. sonnei* scanned (625/3,428) had an SSU5 group P-P. Genotyping of the 625 SSU5-positive *S. sonnei* whole genomes revealed that those with P-Ps primarily belonged to Lineage III (n=623, 99.7%), in particular the ciprofloxacin-resistant sub-lineage 3.6.1.1.2 CipR.MSM5 (n=510, 81.6%), which is extensively drug-resistant (XDR) and causes a high burden of disease within the MSM (men who have sex with men) community (40–42).

Of the 497 SSU5 group P-Ps found in *S. enterica*, *S.* Typhi represented 51%: 17% of all *S.* Typhi that were scanned (254/1,517) possessed an SSU5 P-P. Genotyping of the 254 SSU5-positive *S.* Typhi whole genomes showed that 75% (191/254) belonged to sub-lineages 4.3.1.1 and 4.3.1.2, also known as haplotype 58 (H58), which is MDR and globally disseminated. The SSU5 group P-Ps also had a wider distribution within *S. enterica* than the P1-like P-Ps; outside of *S.* Typhi there were 243 P-Ps present across 59 other serovars and spanning four of the sub-species of *S. enterica*.

In the *Escherichia* dataset, as seen with the P1 group, SSU5 P-Ps were particularly common in the STEC/ExPEC hybrid CC165 (61% of isolates) and were again associated with the MDR, hypervirulent ST301 sub-lineage. We also noted a high frequency (16%) among the CC131 isolates, which are scarce in our collection as this complex is typically an extra-intestinal pathogenic *E. coli* (ExPEC) that causes urinary infections and bacteraemia rather than gastrointestinal illness. In contrast, the prevalence among other important CCs and serovars within our collection – particularly those aforementioned groups with higher frequency of P1-like P-Ps – was far lower; for example, 23 in *S.* Typhimurium (0.26%; n=21 in eBG1 and n=2 in eBG138), six in *E. coli* CC11 (0.14%) and seven in CC29 (0.65%).

The SSU5 supergroup has been shown to contain several defined sub-groups such as SSU5_pHCM2 and pSLy3 (5,15), and so it was important to compare the population of P-Ps analysed in the current study. We downloaded related, publicly available genomes and combined these with the extracted P-P contigs from our collection to examine the overall population structure (Fig. 4). This SSU5 community tree comprised 11 distinct clades (A-K), eight of which contained P-Ps from our UK collection. The host species displayed a clear separation, with clades C-F and H-I containing all P-P contigs from the *S. enterica* isolates and clades J-K containing all P-P contigs from the *Escherichia* and *Shigella* isolates; apart from two *E. coli* CC11 elements in clade D that were closely related to one from *S.* Mikawasima (Fig. S9).

**Figure 4.**
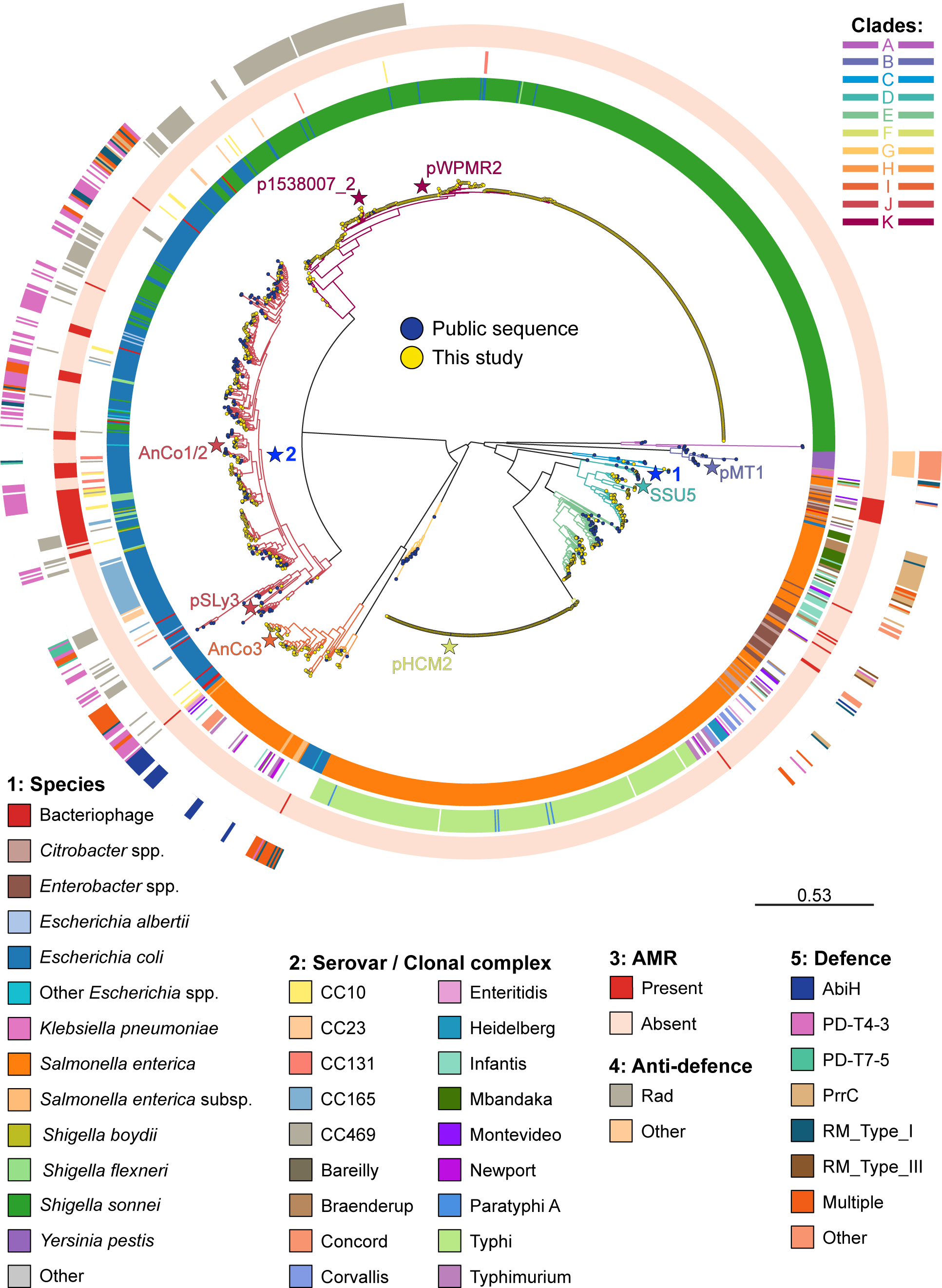
Midpoint-rooted tree showing the population structure of the SSU5 P-P group (n=1529 total), including public sequences, with metadata of bacterial host strain where available. Main serovars (*Salmonella*) and clonal complexes (*Escherichia*) are shown, the location of references and sequences of interest are labelled, and the two AMR clades (Fig. 5) are numbered and highlighted with a blue star.

We noted that clade F contained highly conserved P-Ps deriving only from *S.* Typhi (248/252) or *S.* Paratyphi A (4/252) isolates, and that these were identical or nearly identical to the well-characterised pHCM2 (23). Likewise, clade K contained highly conserved P-P sequences that were largely from *S. sonnei* (540/574) and were identical to, or a close match with, the recently described pWPMR2 (43), which is a P-P associated with numerous outbreaks of XDR shigellosis among the MSM community.

Supporting this, within clade K we also found p1538007_02, which similarly derives from an XDR, UK MSM-associated isolate of *S. sonnei* (42). Similarly, a sub-clade within clade J contains P-Ps only from CC165. These examples are suggestive of sustained vertical inheritance within the respective ‘primary host’ populations. As was seen in the P1/D6 P-P tree, we also detected numerous examples of apparent horizontal transmission occurring between various species, serovars and CCs, and again spanning time and geography (Fig. S9).

After scanning the SSU5 group dataset for resistance factors, we detected 45 P-P contigs with AMR genes (Table 2). Of these, 37 had *bla*_CTX-M-15_ carried on an IS*Ec9* element, which was located at different sites upon the P-Ps themselves though noticeably excluding the region containing the conserved phage structural (capsid, portal, tail, fibre) genes (Fig. 5A). In three *S. enterica* isolates from different serovars we found SSU5-like P-Ps (clade D) that were related to elements from *Citrobacter* and *Enterobacter* and carried multiple AMR genes on Class I integrons (AMR Clade 1 in Fig. 4, Fig. 5B). We also detected *bla*_TEM_ alleles on Tn*3*-like unit transposons in an *E. coli* CC40, an *E. albertii* and an *S.* Typhimurium eBG1, all inserted near tRNA loci (Fig. 5C). Overall, the P-Ps carrying AMR genes came from a diverse range of species and STs/serovars and were located in multiple clades across the overall community (Fig. 4). Notably however, no AMR genes were detected in P-Ps from either of clades F (pHCM2) or K (pWPMR2), despite their frequency among our collection.

**Figure 5.**
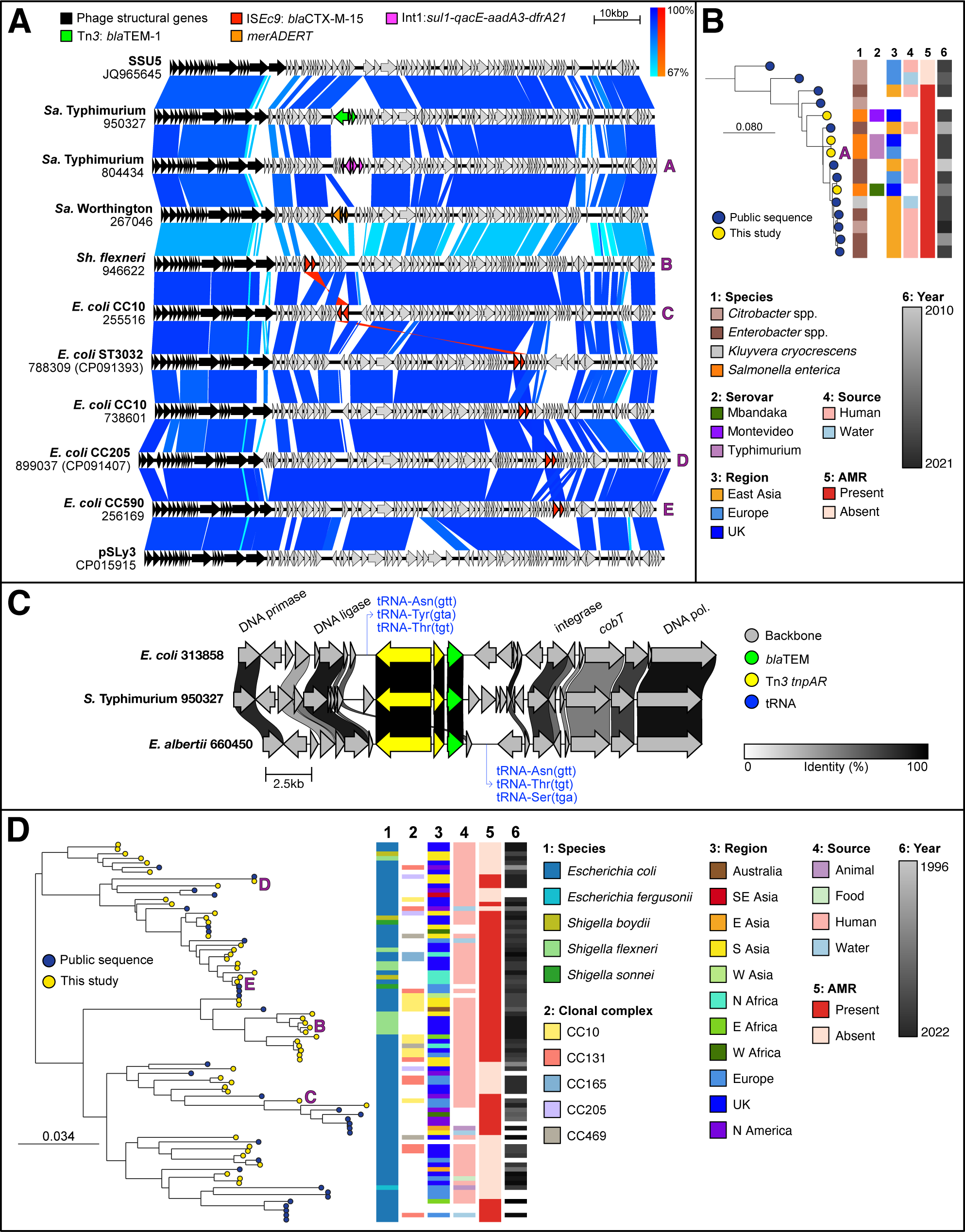
Antimicrobial resistance (AMR) in phage-plasmids (P-Ps) belonging to the SSU5 supergroup. **A)** Nucleotide sequence relatedness between selected P-Ps from this study that were found to carry AMR genes, alongside the reference P-P genomes SSU5 (top) and pSLy3 (bottom), showing differing locations of AMR gene cassettes and overall sequence diversity within the SSU5 supergroup. **B)** Detailed view of AMR clade 1 (within clade C), midpoint-rooted tree showing *S. enterica* P-Ps carrying Class I integrons with multiple AMR genes, and their close relationship to public sequences from other species. **C)** Comparison of the locations and surrounding features of the *Tn3* transposons carrying *bla*_TEM_ genes in three P-Ps from different species. **D)** Detailed view of AMR clade 2 (within clade J), showing P-Ps carrying IS*Ec9* elements with the *bla*_CTX-M-15_ gene, and their relationship to public sequences. Metadata for public sequences is displayed where available; region is based on travel data for UK isolates (where known) or isolation locations for public sequences.

Of the 39 AMR-containing contigs that were extracted and included in the tree, the majority (79%) were situated in a single phylogenetic group within clade J (pSLy3 subgroup (5)) alongside numerous public sequences (such as pAnCo1 and pAnCo2 (29)) that also carried the *bla*_CTX-M-15_ gene. Within this sub-clade (AMR clade 2 in Fig. 4, detailed in Fig. 5D) we observed several further examples of close relationships between P-Ps from this study and geographically distributed elements in the public domain. Firstly, three from *E. coli* and *S. boydii* (received between 2016-2020, all reporting travel to Egypt) were identical to P-Ps from *S. sonnei* and *E. coli* CC131 (2017-2018, Belgium and Sweden respectively, with no reported travel) (44,45). Elements from two *E. coli* isolates with reported travel to India and Pakistan between 2018-2019 (37), alongside one each from *S. sonnei* and *S. boydii* with unknown travel, were highly similar to a P-P from another diarrhoeagenic *E. coli* with reported travel to India in 2018 (46), and one sampled from a chronic wound of a patient in Ghana (47).

Another example within AMR clade 2 covers P-Ps from two *E. coli* isolates with no reported travel, which were related to P-Ps from *E. coli* in sewage in India (48), ExPEC from a dog in Thailand (49), diarrhoeagenic *E. coli* sampled from soldiers deployed in Mali (50), and environmental *E. coli* from wildlife in the USA from 2013-2014 (29). Lastly, the sequences from two *E. coli* CC10 isolates, deriving from patients with travel to India in 2016, were >99% identical to P-Ps isolated between 2014-2016 in Lebanon and Australia, also both belonging to *E. coli* CC10 hosts sampled from human infections (51,52).

Regarding anti-defence systems (Table 3), in contrast to the P1/D6 P-Ps there was a notably lower frequency and diversity in the SSU5 group: 16% of P-P sequences from *Escherichia* and 22% from *Shigella* had the Rad anti-Retron system, while we found zero systems in the *S. enterica* P-Ps. Most of the P-Ps carrying Rad belonged to *S. sonnei* (132/161, 82%) and were located in clades J or K only (Fig. 4). We also observed that defence systems were again less frequent – yet more diverse – in the SSU5-like P-Ps, when compared to the P1 elements. Differences included the presence of AbiH, DS-33, DS-37 and HEC-06 systems only among P-Ps from *S. enterica*; while the P-Ps in clade J that were found in *E. coli* and *Shigella* spp. instead had PrrC, PD-T4-3, PD-T7-5 or RM systems. As with the resistance genes, there were no detected defence systems in any of the pWPMR2-like P-Ps belonging to clade K, nor in the pHCM2-like *S.* Typhi P-Ps in clade F. In general, the defence systems found in the P-Ps belonging to the SSU5 supergroup were not shared with the P1/D6 group, and vice versa, with the exception of the PrrC and RM type III systems (Table 4). Predicted systems particularly included the Lamassu-Fam nuclease system and the DEDDh proteins that are ancillary exonucleases of CRISPR-Cas systems, as well as various putative RM systems (Fig. S8).

### N15 Group

The linear N15-like P-Ps were least commonly found, and most (n=99, 84%) were detected in the genomes of *S. sonnei* isolates. Based on SNP typing data of the host isolate genomes, these were distributed sporadically rather than being associated with particular clusters, apart from n=18 in isolates from a single MSM-related outbreak cluster (Son26, lineage 3.7.25 MSM4) that occurred between 2017-2018. In *Escherichia* spp. (n=14) and *S. enterica* (n=5) the N15-like P-Ps were also largely sporadically acquired: these were located over twelve different STs and five serovars, respectively.

As the N15-like P-Ps were largely found in *S. sonnei* and *S. flexneri*, which are notoriously difficult to assemble from short-read sequencing data, we were limited to a total of only seven contigs >40 kbp for comparative analysis; these however spanned five species. The seven N15-like contigs showed a conserved layout with differences occurring mostly in the phage tail encoding genes (Fig. S10) and a mosaic relationship to other phages and linear P-Ps. The region containing phage structural genes (also referred to as the structural gene cassette, SGC (18)) shared considerable identity with *Escherichia* phage Lambda and other lambdoidal phages. Meanwhile, there were similarities between the plasmid region (containing the *repA* replication gene, *sopAB* partitioning system and *telN* protelomerase) and P-P elements from *Klebsiella* such as phiKO2 (19) and the recently described NAR688 that is widespread in *K. pneumoniae* (18).

With our limited number of full-length contigs, we detected only one instance of an almost identical P-P being present in different hosts: elements from an *E. marmotae* and an *E. coli* ST139. Among public genomes, we noted similarity to p3 of OXEC-387, found in an isolate from a UK bacteraemia patient in 2017 (53), and also to an element from an isolate of *S. sonnei* collected in France in 2014 (18,54). No known or putative AMR or virulence genes were found in any of the N15-like P-P sequences, nor any complete defence systems, though there were predicted DEDDh proteins in all sequences and putative Type II RM systems in six of the seven sequences. Furthermore, we did not detect any homologues of the recently characterised telocin that is associated with the NAR688-like telomere phages (18).

## Discussion

This is the first systematic study exploring the presence of phage-plasmids in a large cohort of clinical Enterobacteriaceae samples and assessing their relevance to public health. It is based on a comprehensive dataset of approximately 67,000 clinical isolates collected at UKHSA from 2016 to 2021 and subsequently submitted to public databases. While there are numerous studies on the diversity of plasmids and phages in Enterobacteriaceae, which include commentary on their impact on human health (53,55–57), there has not yet been any large-scale surveillance of P-Ps that links back to host diversity below species level, and/or incorporates clinical and public health importance. This study expands on previous ones (11,15,26,27), adding a detailed characterisation of genomic features of both P-Ps and their hosts.

### P-Ps are an understudied mobile element in clinically relevant species

There is little evidence that existing bioinformatics tools and public health surveillance pipelines are optimised towards recognition of P-Ps. We note that common plasmid detection and typing tools include P1 (IncY and p0111) and SSU5 (IncFIB) replication genes in their databases (58,59), however other P-P groups (D6 and N15 at least) are not included, despite their apparent importance in human pathogens. For example, D6-like P-Ps are associated with invasive *S.* Typhimurium strains (60), and N15-like P-Ps (or ‘telomere phages’) were recently revealed to be widespread in *Klebsiella* spp. and importantly often possess a bacteriocin that impacts interbacterial competition (18).

Genomic epidemiology studies that perform plasmid surveillance often mis-identify P-Ps only as ‘cryptic plasmids’ or exclude them entirely (5,15,61–63). Our screening shows that while numbers per P-P group may be low, overall totals are considerable, and P-Ps are found in ∼8% of *Escherichia*, ∼10% of *Shigella* and ∼1/50 *Salmonella* isolates that have caused human gastrointestinal disease. This has clear implications towards public health, and it is important to gain a better understanding of their distribution in pathogenic species.

### The relation between P-P group and host is a complex one

Only a comprehensive study of P-Ps can successfully elucidate the relationship between P-P types and their hosts, due to the complex interplay between horizontal and vertical transmission. The scope of our study allowed us to discover that some types of P-Ps are prevalent in specific host species, lineages and/or clades, while others are rarer, dispersed across the various populations, and therefore likely sporadically acquired through HGT.

For instance, we find pHCM2-like P-Ps in almost a fifth of *S.* Typhi, which is a similar prevalence to other studies (64), but also identified that near-identical elements are present at a lower rate in *S.* Paratyphi A, and that more diverse SSU5-like P-Ps are present in numerous other clinically relevant serovars, including *S.* Typhimurium, *S.* Newport and *S.* Infantis. We confirm the reports of pWPMR2-like P-Ps in a considerable percentage of *S. sonnei* (43) and show that this P-P is also present in *E. coli* complexes such as CC131, CC23 and CC73. Furthermore, when considering this study in the context of our previous report on P1 P-Ps in *S. enterica* (11), we see that these elements are in fact much more prevalent among diarrhoeagenic *E. coli* and *E. albertii* and so are apparently only sporadically acquired by *S. enterica*.

Our study identified several instances that are clearly relevant to public health; specific combinations of P-P group and host CC/serovar/lineage, many of which are human-adapted pathogens lacking animal reservoirs (65–68), and that have links to more serious disease and/or are resistant to multiple antibiotics. For example, *E. coli* ST301 (CC165) is an emerging MDR STEC/ExPEC hybrid clone that is highly virulent and can cause severe, systemic disease. The ST301 pSLy3-like P-P has been previously noted but referred to only as ‘cryptic plasmid’ pR444_B (32,33,63), and we additionally showed that almost a quarter of the CC165 genomes in the UKHSA archive have a second, P1-like P-P. Furthermore, we identified a number of isolates belonging to the epidemic MDR ExPEC CC131 complex that also carry pSLy3-like P-Ps, and highlighted the particular association of pHCM2 (also described as cryptic (23)) with *S.* Typhi lineage 4.3.1 (H58), which is globally spread and associated with high AMR gene carriage (69). Lastly, we emphasised the widespread prevalence of a highly conserved P-P among *S. sonnei* MSM lineages (43) – again known for being XDR and responsible for a large proportion of the domestic shigellosis cases seen in the UK (40–42,70,71).

An earlier study of invasive *S.* Typhimurium ST213 highlighted a link between D6-like P-Ps and MDR IncC plasmid presence (34,60). We note that the aforementioned *E. coli* ST301 and CC131, and *S.* Typhi 4.3.1 (H58), are also commonly invasive in humans, and importantly that these pathogens alongside *S. sonnei* 3.6.1 are all strongly associated with carriage of resistance plasmids (32,63,72–74): this pervasive connection between P-Ps and plasmids in prominent human pathogens merits further investigation. In stark contrast to the above, we also found notable under-representations in our dataset: P-Ps were only found in eight *S.* Enteritidis isolates, despite this serovar contributing to more than a quarter of the UK *S. enterica* collection; and, when compared to *S. sonnei*, the other three species of *Shigella* have considerably lower P-P carriage.

### Correlation between P-Ps and their hosts provides genomics insights

Regarding the bacterial chromosomal diversity, the breadth of our study also allowed us to correlate the phylogeny of P-Ps with that of their hosts for the first time, which is much more meaningful than looking at P-Ps in isolation. The prevalence of particular P-P groups among specific STs and SNP clusters suggests likely vertical transmission in several cases. Likewise, the distribution of similar or identical elements across different host lineages and species suggests that horizontal transmission is occurring across species and genera, albeit largely sporadically. This is expected, as P-Ps by their nature can disperse in the environment (as phage particles following lysis) rather than requiring cell-to-cell contact (4). Furthermore, examining the P-Ps in the context of their host bacteria allowed us to look at the instances where isolates carried two P-Ps at once, providing evidence that the various groups are not incompatible: even the close relatives P1 and D6 could be found together, in a few *E. coli* isolates.

### Genes relevant to public health are associated with clades of P-Ps

Another notable finding is our detection of important accessory genes associated with clades of P-Ps, and their association with host and epidemiological metadata. There are numerous studies reporting AMR gene carriage in both the P1 and SSU5 groups within Enterobacteriaceae hosts (9,25–29,31). While some limited further AMR determinants were found in the P1 P-Ps examined in this study, we did identify a globally distributed clade of pSLy3-like (SSU5) P-Ps with carriage of IS*Ec9* containing *bla*_CTX-M-15_, with public sequence metadata indicating links to water and evidence of international spread via travel. This promiscuous AMR element was present across multiple species and CCs and had inserted into a variety of locations within the P-P sequences while avoiding conserved phage genes as previously observed (9). We also detected other types of elements embedded into SSU5 P-Ps, including class I integrons and Tn*3*-like unit transposons, evidencing how P-P backbones can acquire diverse elements that may subsequently impact on clinical disease. It was discovered previously that P1-like P-Ps may transition into plasmids by acquiring genes necessary for conjugation such as *mob* relaxases, and that these novel P1-derived plasmids very often carry AMR genes (6). Our finding of multiple international P-P clades with high frequency of AMR gene carriage, occurring in both the P1 and SSU5 groups, lends further evidence towards certain sub-populations of P-Ps evolving towards being established AMR vectors – with or without the gain of plasmid mobility genes.

Furthermore, we report for the first time the presence of the cytotoxic factor Cnf3 linked to a large clade of P1-like P-Ps that are circulating amongst the STEC population, in particular CC29 (O26:H11) and some CC11 (O157:H7). P-Ps are already known to be important for pathogenicity in some species, such as the pMT1-encoded murine toxin Ymt in *Y. pestis* that is crucial for transmission from fleas to humans (75). More recently, a P1-like P-P with a large virulence region containing *stb* (heat stable enterotoxin II) and *eltAB* (heat labile enterotoxin) was found in porcine enterotoxigenic *E. coli* O157 (76). Cnf3 shares considerable sequence homology with Cnf1, an important virulence factor in ExPEC strains, suggesting that it may have similar effects on pathogenesis such as increased inflammation or impacting immune cell function (36). Further studies are required to investigate the effect the P1-encoded *cnf3* may have on virulence in pathovars of *E. coli* such as STEC, but our finding is important considering that CC29 strains are often linked with more serious disease (such as haemolytic uraemic syndrome, HUS) in the UK (77,78).

In addition, we screened the P-P contigs for anti-defence systems that protect from host killing mechanisms. Certain systems are very well characterised in P-Ps, such as the P1 defence against restriction system (38,39), and genes from various anti-defence systems have been found in the pan-genomes of most P-P groups (5). As expected, we found anti-defence systems across the majority of P-Ps in our dataset; particularly anti-RM and anti-Thoeris systems in the P1/D6 group, and the Rad anti-Retron system in the subset of SSU5 sequences linked to XDR *S. sonnei* (43,79). These likely contribute to the stability of P-Ps in their hosts, increasing their chances of inheritance. Alongside anti-defences providing protection from their hosts, we also found various defence (i.e. anti-phage) systems. These included RM Type III among the P1-like P-Ps – particularly in *S.* Typhimurium – and putative RM systems in many more P-Ps from all groups, as well as numerous other abortive infection systems that are already known to be present widely in the Enterobacteriaceae (80). These defence systems may therefore be important for competition with other mobile elements or for preventing lytic phages from destroying their common host.

### Conclusions

There is an increasing awareness of the importance of mobile and extrachromosomal elements to pathogen evolution and adaptation. An increasing number of studies are being published that focus on these mechanisms, either in the clinic or in environmental or other One Health settings. This is the first comprehensive study of a large dataset relevant to public health analysing together the genetic diversity of P-Ps as well as their hosts; shedding light on their co-evolution and interaction. This work demonstrates that P-Ps merit attention as their own category of mobile element, not merely as a by-product of broader studies of plasmids or phages. We believe that this work represents a solid foundation, paving the way for further studies, perhaps of more diverse genome collections that consider other pathovars of *E. coli*, commensal isolates, and those from non-clinical sources.

## Materials and Methods

### Isolates and sequences

*Salmonella* spp., O157 STEC and *Shigella* spp. isolates, or presumptive non-O157 STEC faecal specimens, from hospital laboratories are referred to the gastrointestinal bacteria reference unit (GBRU) at the UK Health Security Agency (UKHSA, formerly Public Health England) for confirmation by PCR, culture and typing, as previously described (81–84). Since 2016, all isolates have undergone routine, short-read whole genome sequencing, and the genomes are regularly uploaded to the NCBI Short Read Archive (SRA) under BioProjects PRJNA315192 (*Escherichia* and *Shigella* spp.) or PRJNA248792 (*Salmonella* spp.). The *E. albertii* collection was additionally supplemented with genomes from a research project (85).

We retrieved the entire dataset available as of 31/12/2021, totalling n=47,495 *Salmonella*, n=12,179 *Escherichia* and 7,182 *Shigella* genomes (Table S1) and the accompanying metadata (Table S2) including strain ID, collection date, serotype/serovar and sequence type (originally derived using MOST (86)). Isolates were assigned to eBURST Groups (eBG, *Salmonella*) or clonal complexes (CC, *Escherichia* and *Shigella*) using the latest available MLST profiles and eBG/CC designations from BIGSdb and Enterobase (87,88). SNP clusters were based on the internal SNP addresses for each isolate using SnapperDB v0.2.9 (89), which represent the relationship between isolates using hierarchical single linkage clustering of pairwise SNP distances at several thresholds (250, 100, 50, 25, 10, 5 and 0 SNPs, respectively). The SNP addresses were simplified to highlight isolates that belong to clusters at the 10 SNP threshold and anonymised via lettering and numbering (e.g. cc11a, cc11b, typhi1, typhi2).

### Long-read sequencing and data processing

The four isolates that had undergone long-read sequencing as part of unrelated projects were sequenced and processed at separate times, and so different processing pipelines were used. For all isolates, DNA was extracted using the Fire Monkey DNA extraction kit (Revolugen) following the manufacturer’s instructions and quantified using a Qubit instrument with the HS dsDNA assay kit (both Thermo Fisher Scientific).

The rapid barcoding kit SQK-RBK004 (Oxford Nanopore Technologies, ONT) was used for library preparation, and prepared libraries were loaded into a FLO-MIN106 R9.4.1D flow cell (ONT) and sequenced using the MinION (ONT) for 48h. Data produced in a raw FAST5 format were base called and de-multiplexed using Guppy v3.2.10 (ONT) using the FAST protocol into FASTQ format and de-multiplexed into each samples’ respective barcode. Nanopore reads were trimmed and filtered using Porechop v0.2.4 and Filtlong v0.2.0 respectively, as previously described (90).

Isolates 671948 and 663175 were draft assembled using Flye v2.6 (91); 1162337 and 1162282 were draft assembled using Flye v2.8 (91). Correction of the draft assemblies occurred in a three-step process. For 671948 and 663175: Nanopolish v0.11.1 (92) followed by Pilon v1.23 (93) and Racon v1.2.1 (94). For 1162337 and 1162282: Nanopolish v0.11.3 followed by Pilon v1.24 and Racon v1.4.20. For all isolates, Pilon and Racon were run with the corresponding Illumina short reads for each genome, as previously described (90). *Shigella sonnei* isolate 1538007 was sequenced and processed as detailed in previous studies (42,71).

### Phage-plasmid screening

Screening for phage-plasmid (P-P) presence in the collection of isolate genomes was carried out as previously described (11). Briefly, reads in FASTQ format were mapped against selected genes for each P-P group (P1: *repL*, *repA-IncY, repA-p0111*; D6: *repL*, *repB*; SSU5: *repB*, *parB*; N15: *telN*, *repB*), with a quality cutoff of at least 1 gene at >10 coverage (i.e. ignoring genomes that matched to P1_*repA*-IncY but not P1_*repL*, to eliminate IncY plasmids), giving a total of 2,701 isolates. FASTQ files for these isolates were downloaded from BioProjects and PRJNA248792. Megahit (95), as part of the Shovill (96) package, was used to perform *de novo* genome assembly (both with default parameters). At this stage, duplicate isolates (n=51), contaminated or mixed genomes (n=51) and false positives (assemblies found to be lacking P-Ps, n=15) were removed, leaving a total of 2,584 assembled genomes containing 2,641 P-P sequences to be analysed further (Table S3). Interestingly, of the 15 false positive genomes, 13 belonged to *S.* Agona and instead appeared to have a chromosomally integrated plasmid with sequence matching the *repB* and *parA* region of SSU5 as well as the P1 p0111 *repA* gene, but lacking phage structural genes, which explained the scanning results and confirmed their exclusion from the P-P dataset (data not shown).

### Phage-plasmid genotyping

To extract and compare P-P sequences, we re-screened the n=2,584 assembled genomes for P1 and SSU5 contigs using a custom script to query the PlasmidFinder database (58), and for D6 and N15 contigs using a BLASTn approach (as these are not covered by PlasmidFinder) and extracted the respective contig that contained the *rep* gene. For assemblies for which we were unable to extract P-P contigs of suitable length (using cutoffs of 75kb for D6, 85kb for P1, 95kb for SSU5 and 40kb for N15 P-Ps), we supplemented the contig dataset by using LastZ v1.04.41 (97) to align the whole genome assemblies to the P1 (NC_005856), D6 (MF356679), or SSU5 (JQ965645) reference sequences, giving a total of 2304 P-P contigs (224/232 D6, 846/921 P1, 1227/1377 SSU5 and 7/111 N15).

These extracted/aligned P-P contigs were screened for antimicrobial determinants using AMRFinderPlus v3.12.8 (98) (Table S4) and for phage defence and anti-defence systems using DefenseFinder v1.3.0 (99–101) (Table S5). Screening for Cnf3 was performed using tBLASTn using the *cnf3* nucleotide sequence with accession CP114131(50688..53729). Reference sequences and selected P-P contigs of interest were compared and visualised using Easyfig v3.0.0 (102) and Clinker v0.0.30 (103), with JalviewJS v2.11.4.1 (104) utilised for the Cnf3 sequence alignment visualisation.

### Trees and comparative genomics

Due to the large size of the *E. coli* CC11 (Fig. S1) and CC29 (Fig. S2) genome datasets, we included all samples harbouring putative P-Ps and a single representative from each 25-SNP cluster. These genomes (n=1,604 CC11, n=1,558 CC29) were aligned to reference genomes STEC O157:H7 strain Sakai (BA000007, CC11) or STEC O26:H11 (NC_013361, CC29) using BWA-MEM v0.7.12 (105) and Samtools v1.1 (106), before using SnapperDB v0.2.9 (89) to produce a soft-core alignment where any given variant position belonged to a minimum of 80% of genomes. Regions of recombination were removed using Gubbins v3.2.0 (107), and the final alignment was used as input to generate a maximum-likelihood phylogenetic tree using IQ-Tree2 (108). For the *E. coli* CC28/CC131/CC165/ST342 (Fig. S4) and *E. albertii* genomes (Fig. S3), whole genome assemblies were input to ParSNP v1.7.4 to generate core genome alignments and RAxML v8.2.12-derived phylogenetic trees for these groups (default parameters). The population structures of the two P-P-positive isolate datasets (*Escherichia + Shigella*, and *Salmonella*), in the form of minimum spanning trees, were constructed and visualised using GrapeTree v1.5.0 with MLST profiles (using MOST (86), as described above) as input.

For the trees generated from the extracted/aligned P1/D6 or SSU5 P-P contigs, sequences were processed using kpop v1.1.1 (109) with a k-mer size of 10 within the kpop-workflow (https://github.com/ryanmorrison22/kpop-workflow), producing neighbour joining, pseudophylogenetic trees. Trees and linked metadata were visualised using the online Microreact platform (110). Related public P-P sequences (Table S6) were identified by BLASTn of reference elements (P1, D6, SSU5 and pSLy3) against the nr nucleotide database, downloading the top hits from GenBank. For overall taxonomy, selected reference P-P sequences were compared to the Virus-Host DB (111) using the VIPTree platform (112).

Genotyping of *S. sonnei* (n=625) and *S.* Typhi (n=254) isolates with SSU5 P-Ps was carried out using the Mykrobe v0.13.0 (113) built-in panels (59,73,114) with the FASTQ reads downloaded from the SRA (Table S7, Table S8).

## Acknowledgements

Acknowledgements to the clinical microbiology laboratories who processed the original patient samples, and to the reference laboratories within the Gastrointestinal Bacteria Reference Unit for culturing and DNA extraction of isolates; to the Central Sequencing Laboratory for library preparation and whole genome sequencing; and to the Core Bioinformatics and Gastrointestinal Genomic Laboratory Services teams for data processing and upload of genomes to the NCBI SRA. The authors acknowledge Research Computing at the James Hutton Institute for providing computational resources and technical support for the “UK’s Crop Diversity Bioinformatics HPC” (BBSRC grants BB/S019669/1 and BB/X019683/1), use of which has contributed to the results reported within this paper (115). This study was funded by the National Institute for Health Research (NIHR) Health Protection Research Unit (HPRU) in Genomics and Enabling Data at University of Warwick in partnership with the UK Health Security Agency (UKHSA), in collaboration with University of Cambridge and Oxford (PR, XD, CB, CC, RM, MC, SN), and supported by the NIHR HPRU in Healthcare Associated Infections and Antimicrobial Resistance at University of Oxford in partnership with the UKHSA (MB, AL) and the NIHR HPRU in Gastrointestinal Infections at University of Liverpool in partnership with the UKHSA (CJ, DG, ER). The views expressed are those of the authors and not necessarily those of the NIHR, the Department of Health and Social Care or the UKHSA. AL was also supported in part by grant NSF PHY-2309135 to the Kavli Institute for Theoretical Physics (KITP) and the Gordon and Betty Moore Foundation grant no. 2919.02.

## Author contributions

Conceptualisation: SN, PR, CB, MB; Data Curation: PR, CB, MB, DG, ER, AP, AC, CJ, MC; Formal Analysis: PR, CB, MB, AL, CC; Funding Acquisition: XD, PR; Investigation: SN, PR, CB, MB, AL, RM, CC, DG, ER; Methodology: PR, CB, MB, AL, RM, CC, DG; Project Administration: SN, PR, CB, MB; Resources: PR, MB, DG, CJ, MC, DG, AP, AC; Software: PR, MB, AAL, CC, DG; Supervision: SN, PR; Validation: PR, CB, MB, AL; Visualisation: CB, MB, AL, DG; Writing – Original Draft Preparation: SN, PR, CB, MB, AL, CC; Writing – Review & Editing: all authors.

## Supplementary Figures

**Fig. S1.**
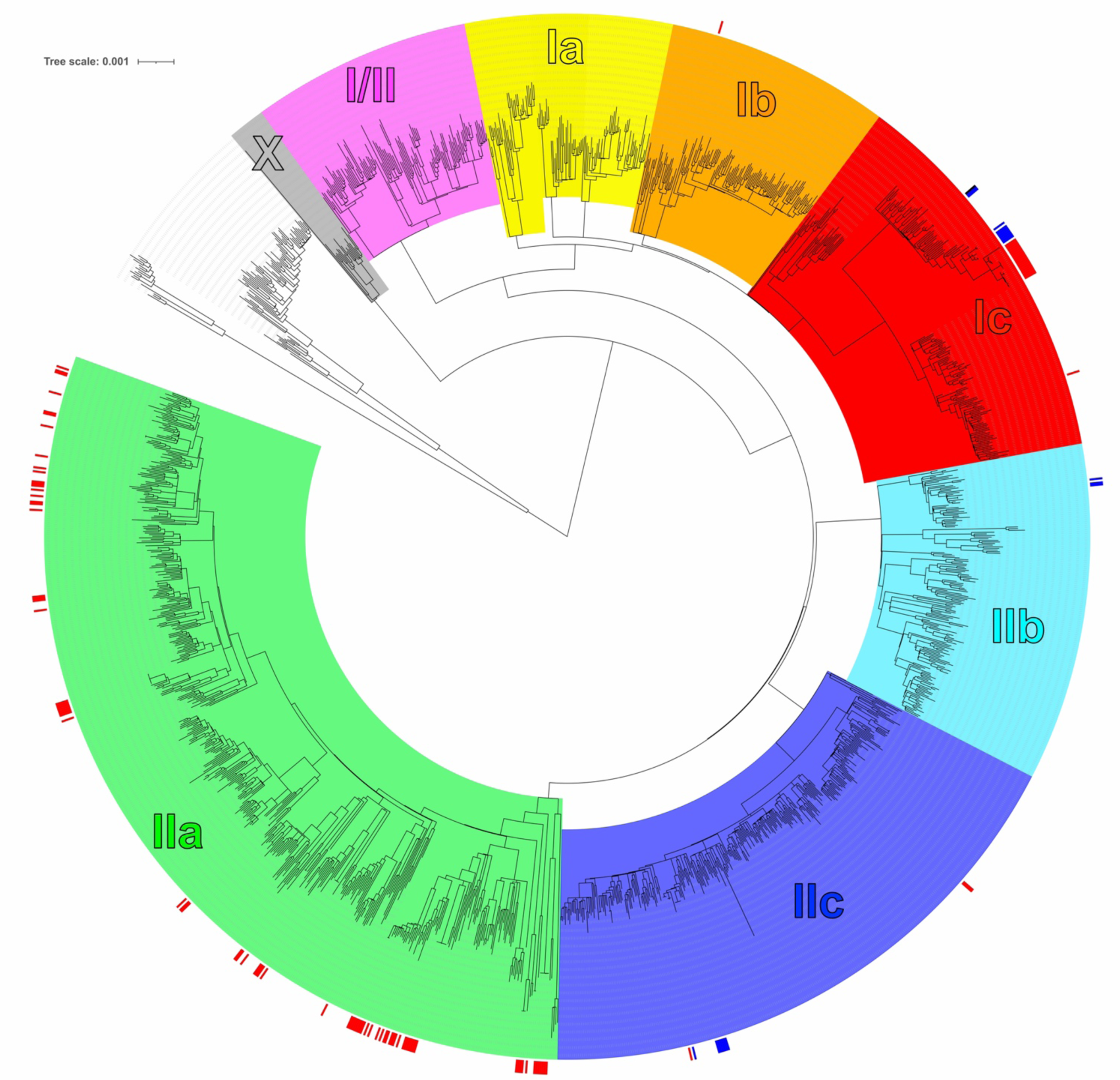
Phage-plasmid distribution in *E. coli* STEC CC11 (O157:H7). Maximum-likelihood phylogenetic tree of n=1,604 isolates with an alignment length of 69,143bp, coloured by lineage (uncoloured = sorbitol fermenting sub-lineage). Outer ring shows P1-like P-P presence, coloured by rep type (p0111=red; IncY=blue).

**Fig. S2.**
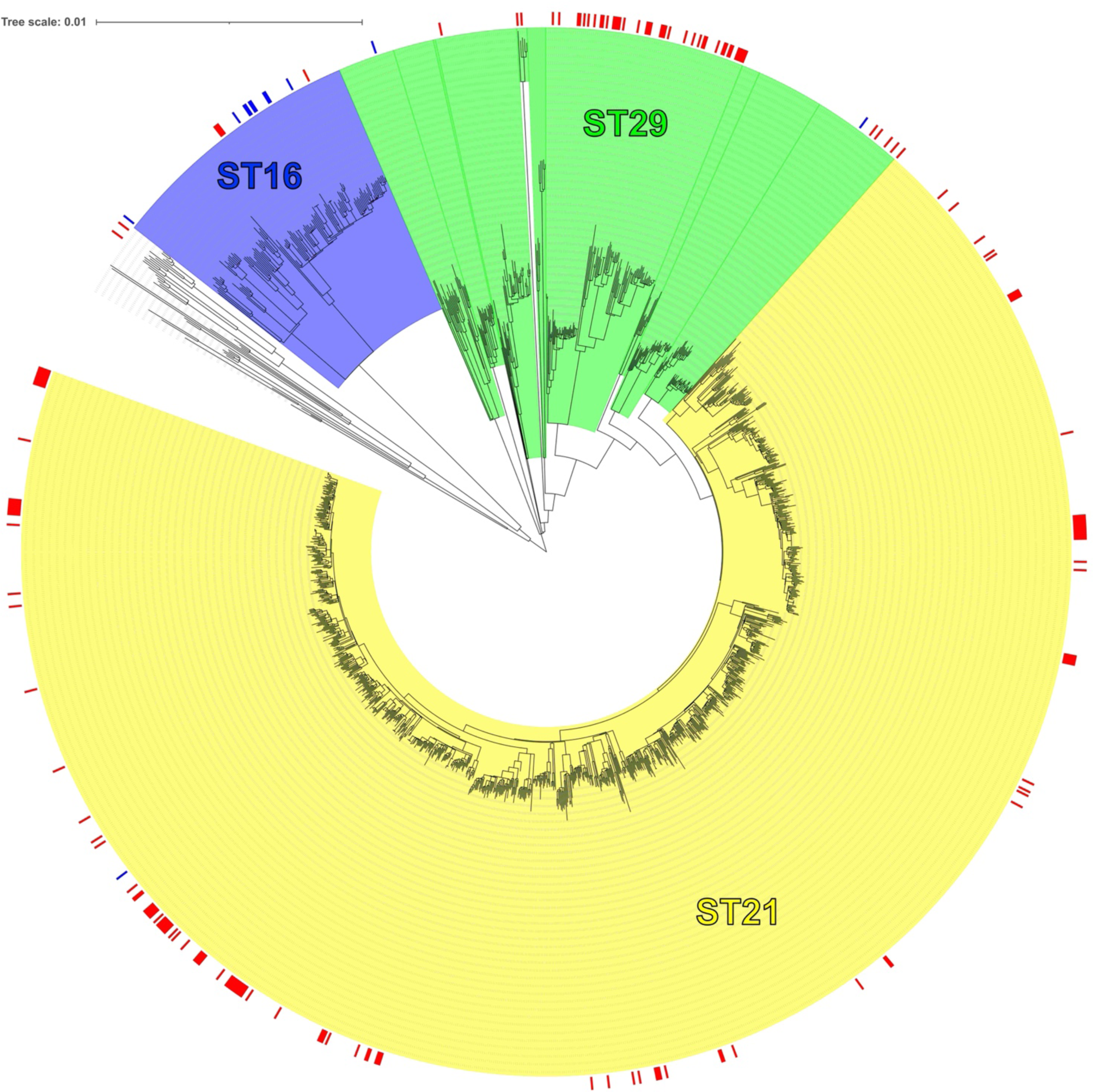
Phage-plasmid distribution in *E. coli* STEC CC29 (O26:H11). Maximum-likelihood phylogenetic tree of n=1,558 isolates with an alignment length of 72,909bp, coloured by sequence type (ST). Outer ring shows P1-like P-P presence, coloured by rep type (p0111=red; IncY=blue).

**Fig. S3.**
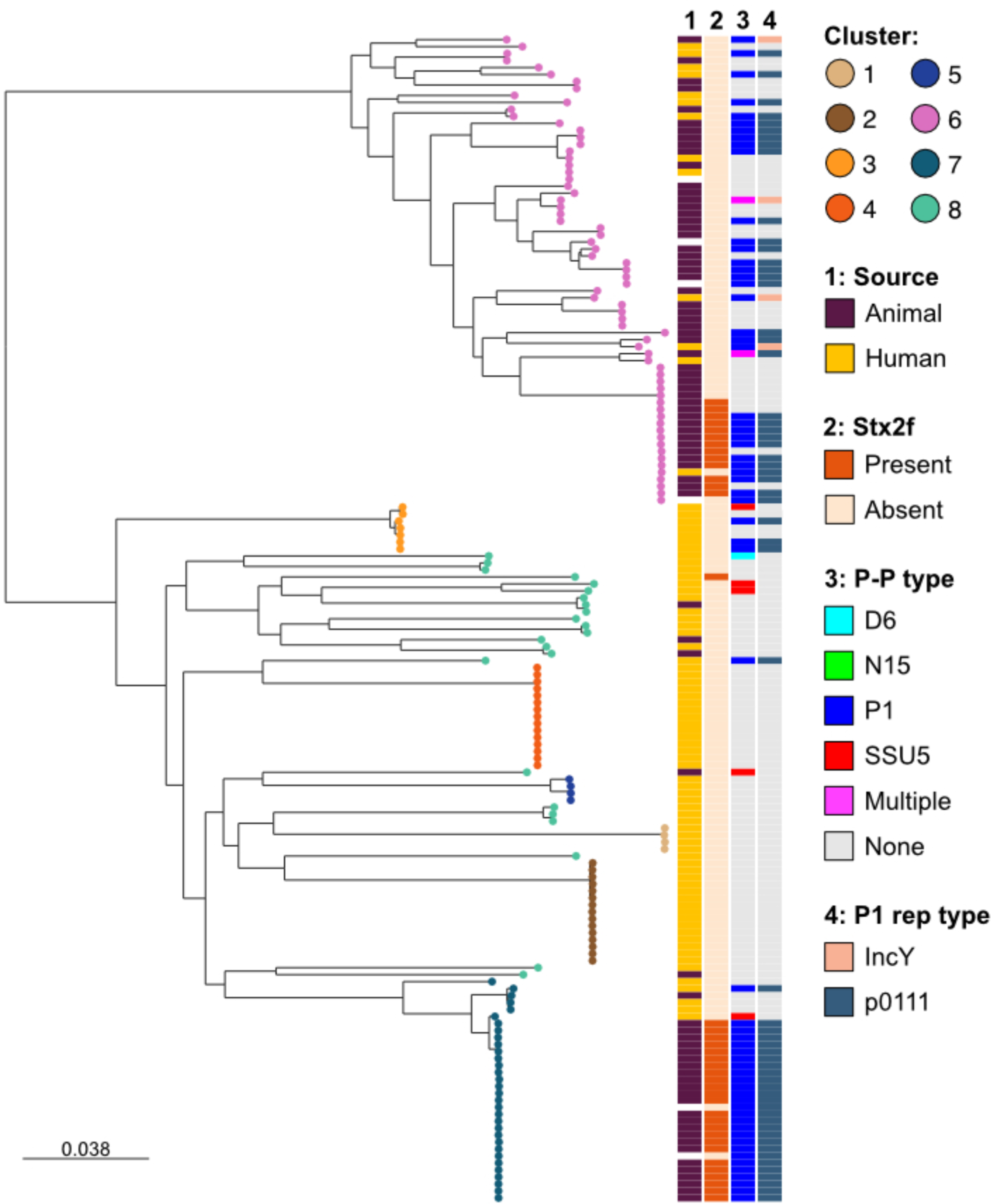
Phage-plasmid distribution in *E. albertii*. Midpoint-rooted, maximum-likelihood phylogenetic tree showing population structure of UK *E. albertii* genomes sequenced between 2016-2021 (n=166), alongside metadata and presence of P-P types. Tips are coloured by clusters characterised previously (85).

**Fig. S4.**
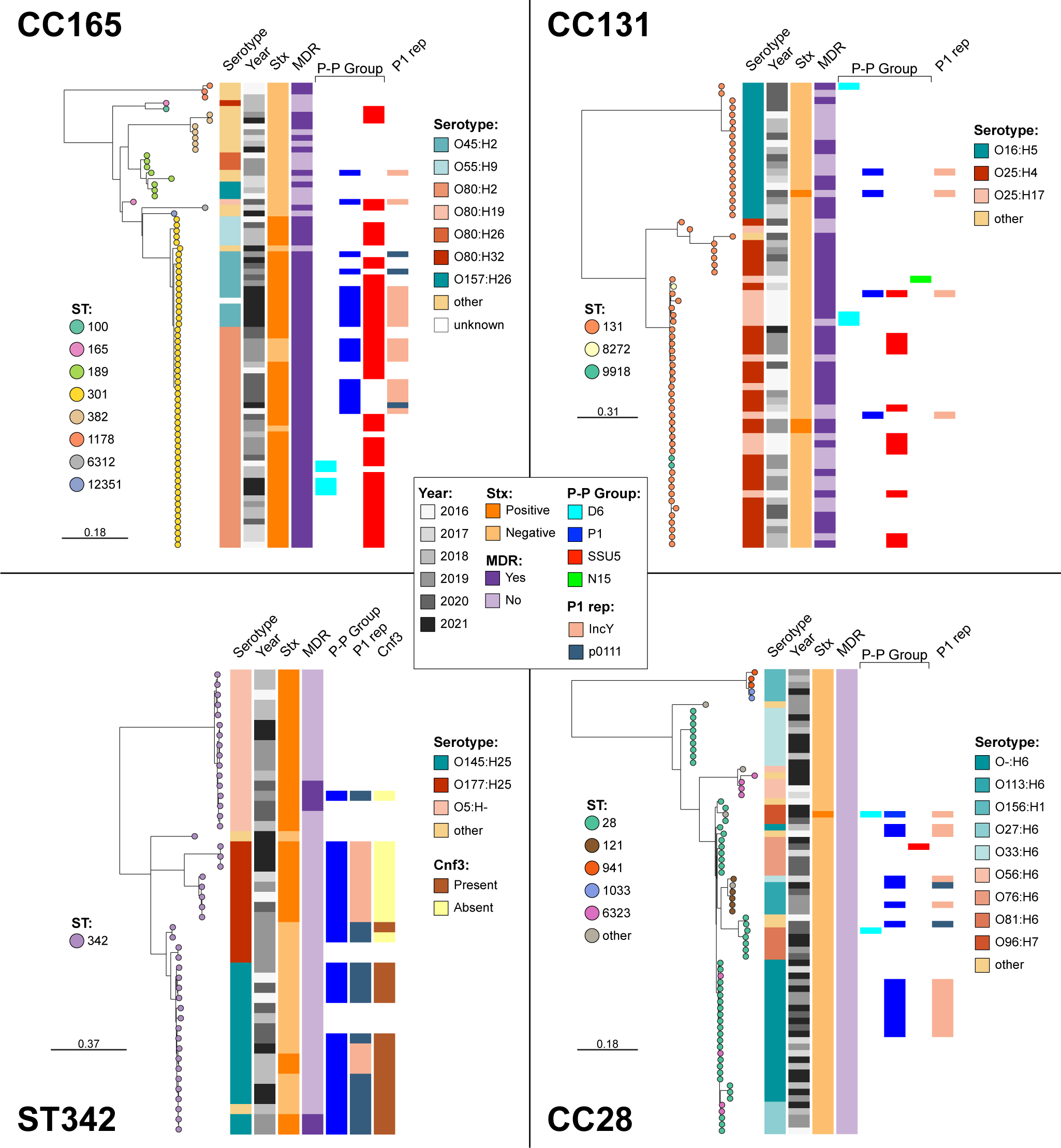
Phage-plasmid distribution in UK *E. coli* sequence types (STs) and clonal complexes (CCs) containing >30 isolates with a P-P frequency >20%. Midpoint-rooted, maximum-likelihood phylogenetic trees showing population structure of: CC165 (70.2% P-P frequency); CC131 (24.6% P-P frequency); ST342 (57.4% P-P frequency); and CC28 (25.7% P-P frequency).

**Fig. S5.**
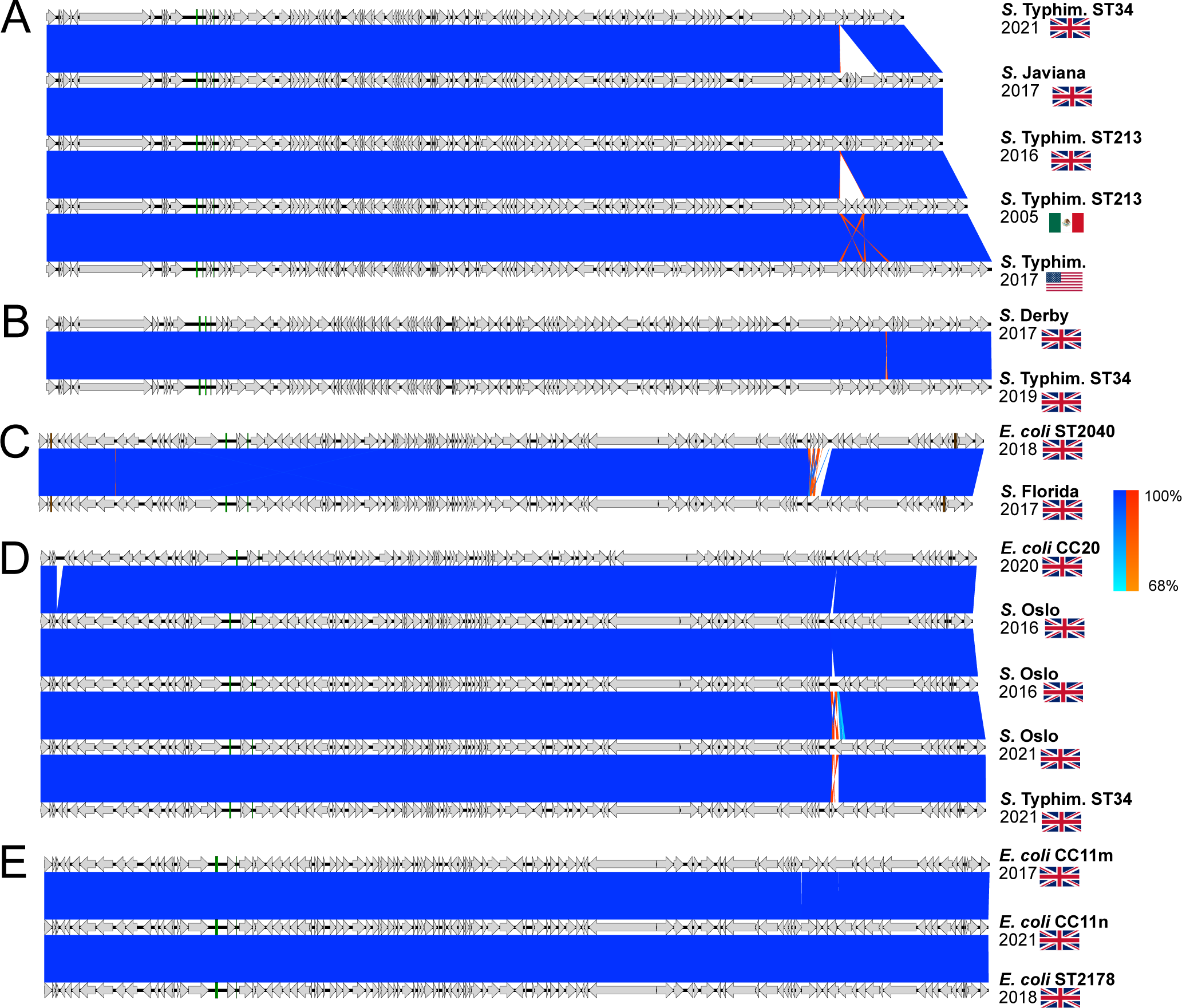
Closely related P1-like and D6-like P-Ps belonging to different hosts, over time and geography. **A,B)** Similarities between D6-like P-Ps in different *S. enterica* serovars. **C,D)** Similarities between P1-like P-Ps in *E. coli* and multiple serovars of *S. enterica*. **E)** Similarities between P1-like P-Ps in *E. coli* CC11 over time and despite host diversity, and ST2178.

**Fig. S6.**
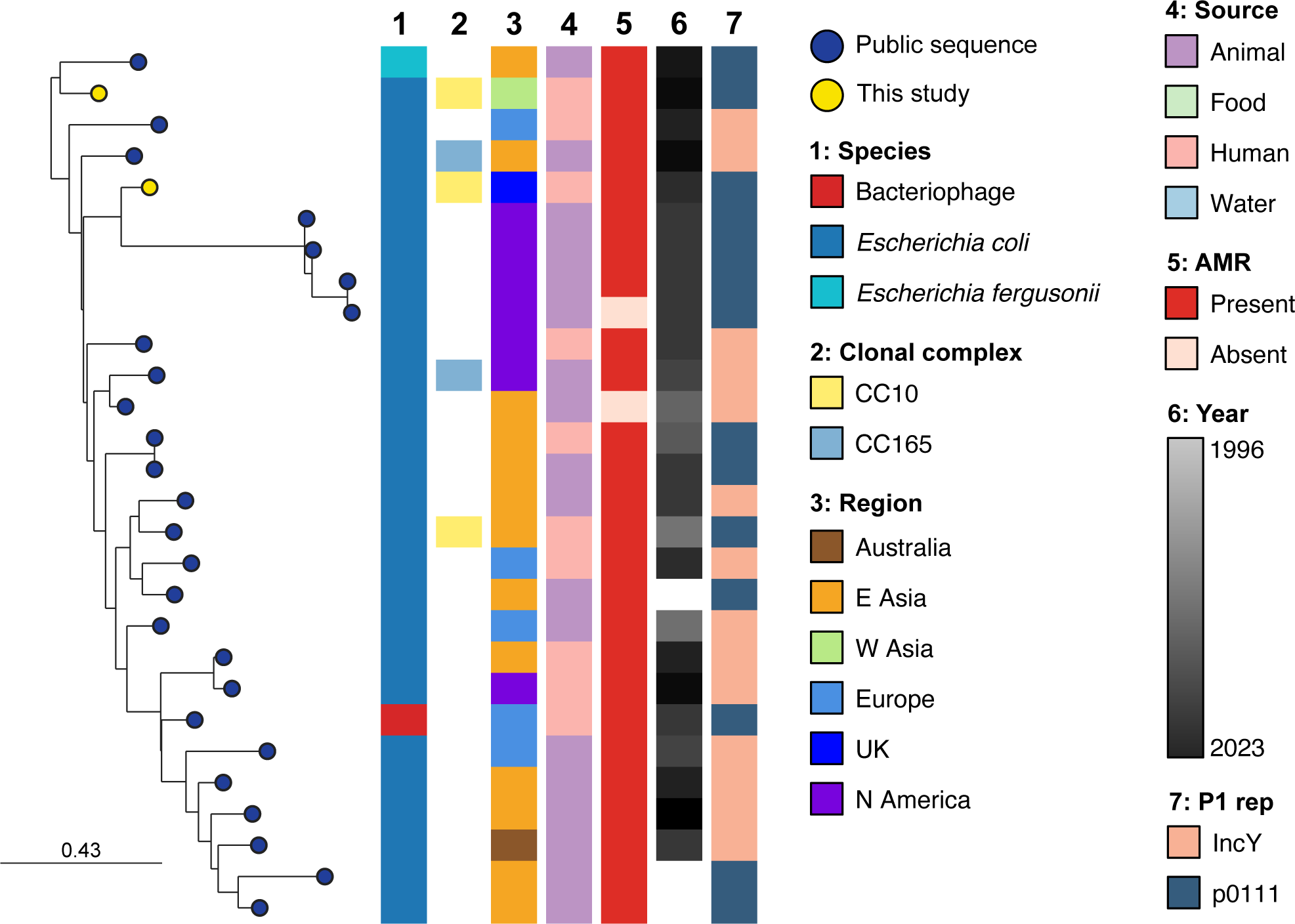
Detailed view of P1/D6 resistance clade, showing P1-like P-Ps carrying various antimicrobial resistance (AMR) genes, and their relationship to public sequences. Metadata for public sequences, including clonal complex (CC), is displayed where available; region is based on travel data for UK isolates (where known) or isolation locations for public sequences.

**Fig. S7.**
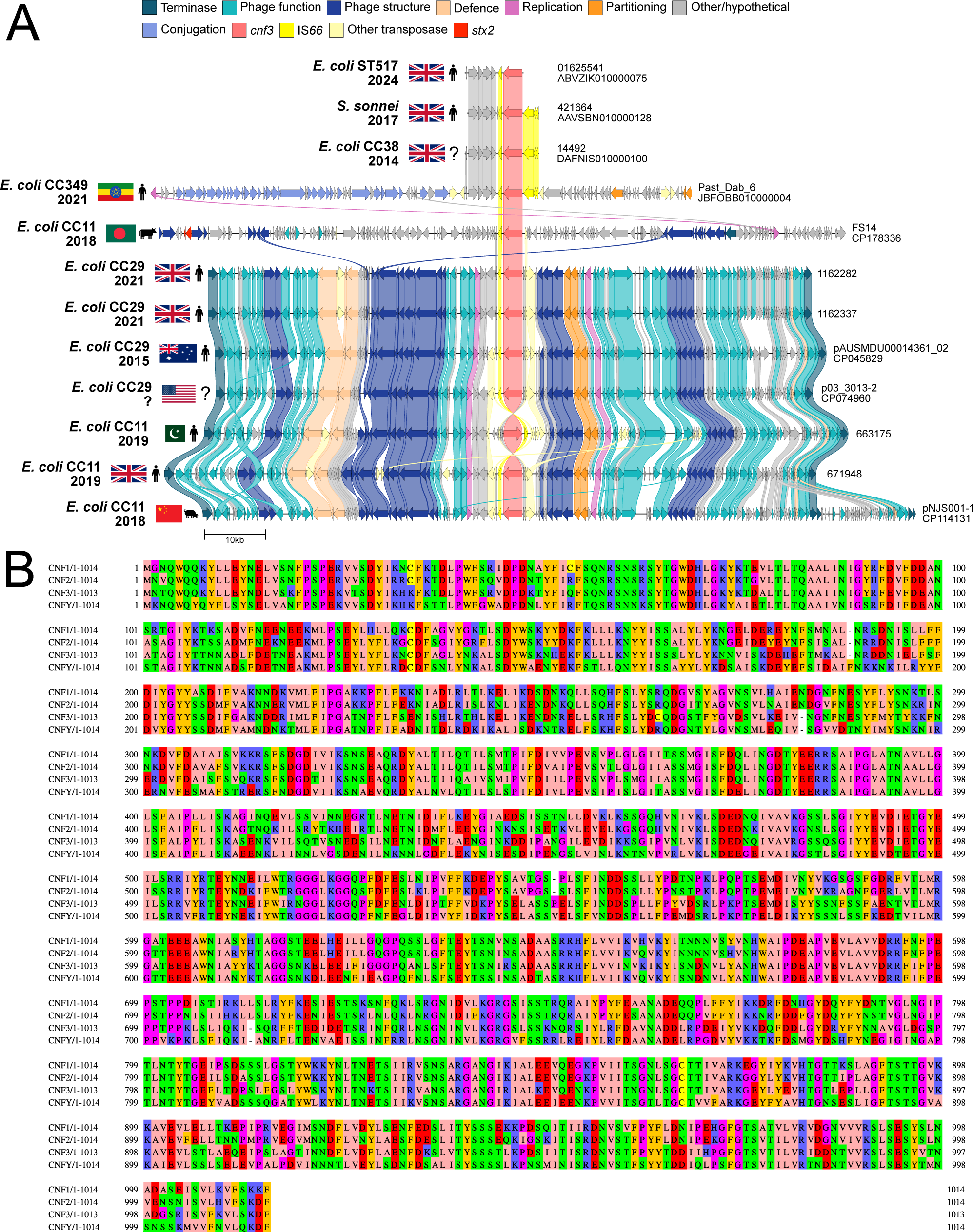
Presence of cytotoxic necrotising factor (CNF3) in P-Ps from Shiga toxin producing *E. coli* (STEC). **A)** Examples of the *cnf3* gene in putative plasmid sequences from UK *E. coli* and *S. sonnei* without P-Ps; in an *E. coli* conjugative plasmid; located near an Stx-converting prophage of STEC CC11 (O157:H7); and among P1-like P-Ps in various STEC strains, with metadata where available. **B)** Visualisation of a Clustal Omega alignment of the Cnf1, Cnf2, Cnf3 and CnfY amino acid sequences.

**Fig. S8.**
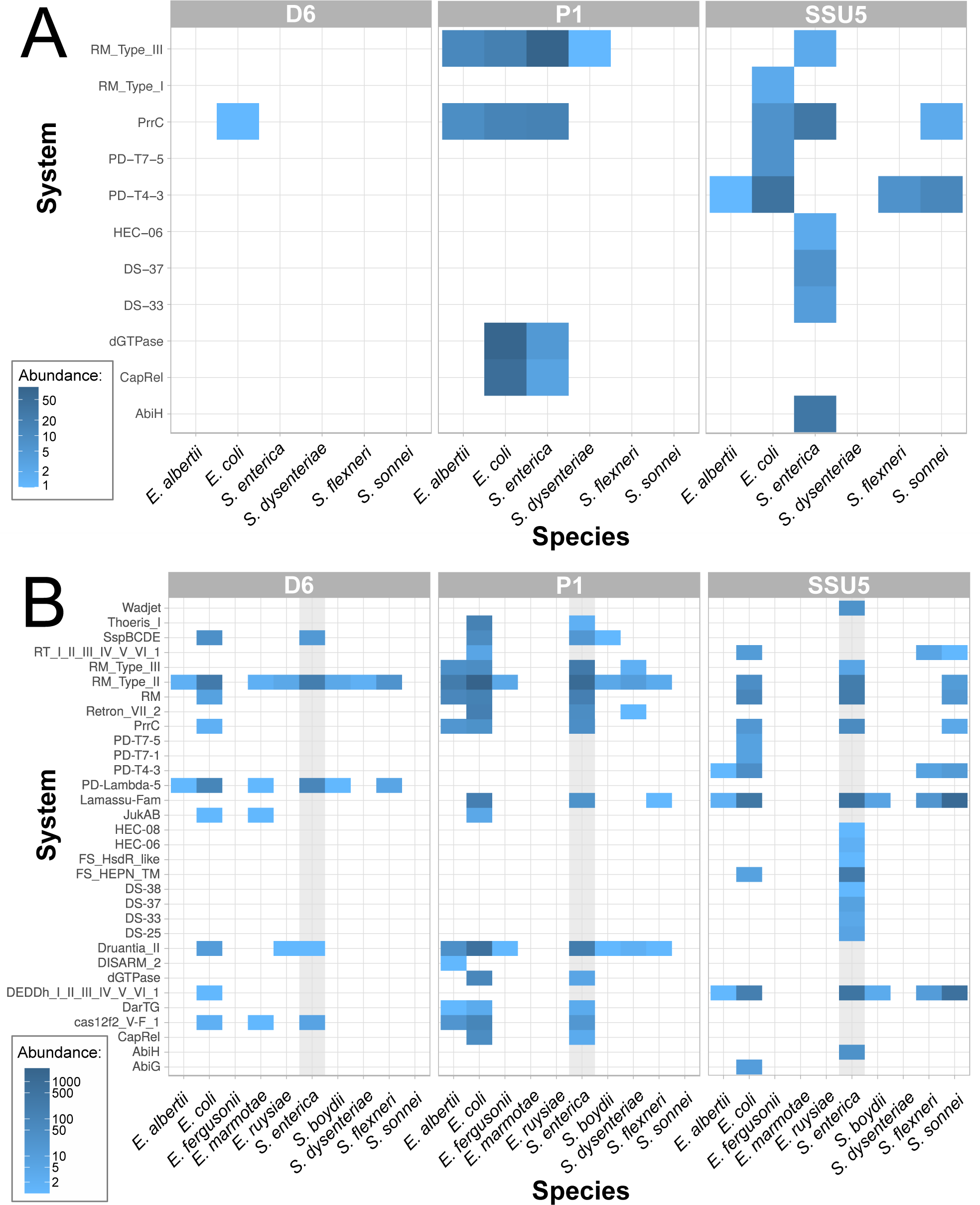
Distribution and frequency of complete (A) and putative (B) defence systems across host species and phage-plasmid (P-P) groups.

**Fig. S9.**
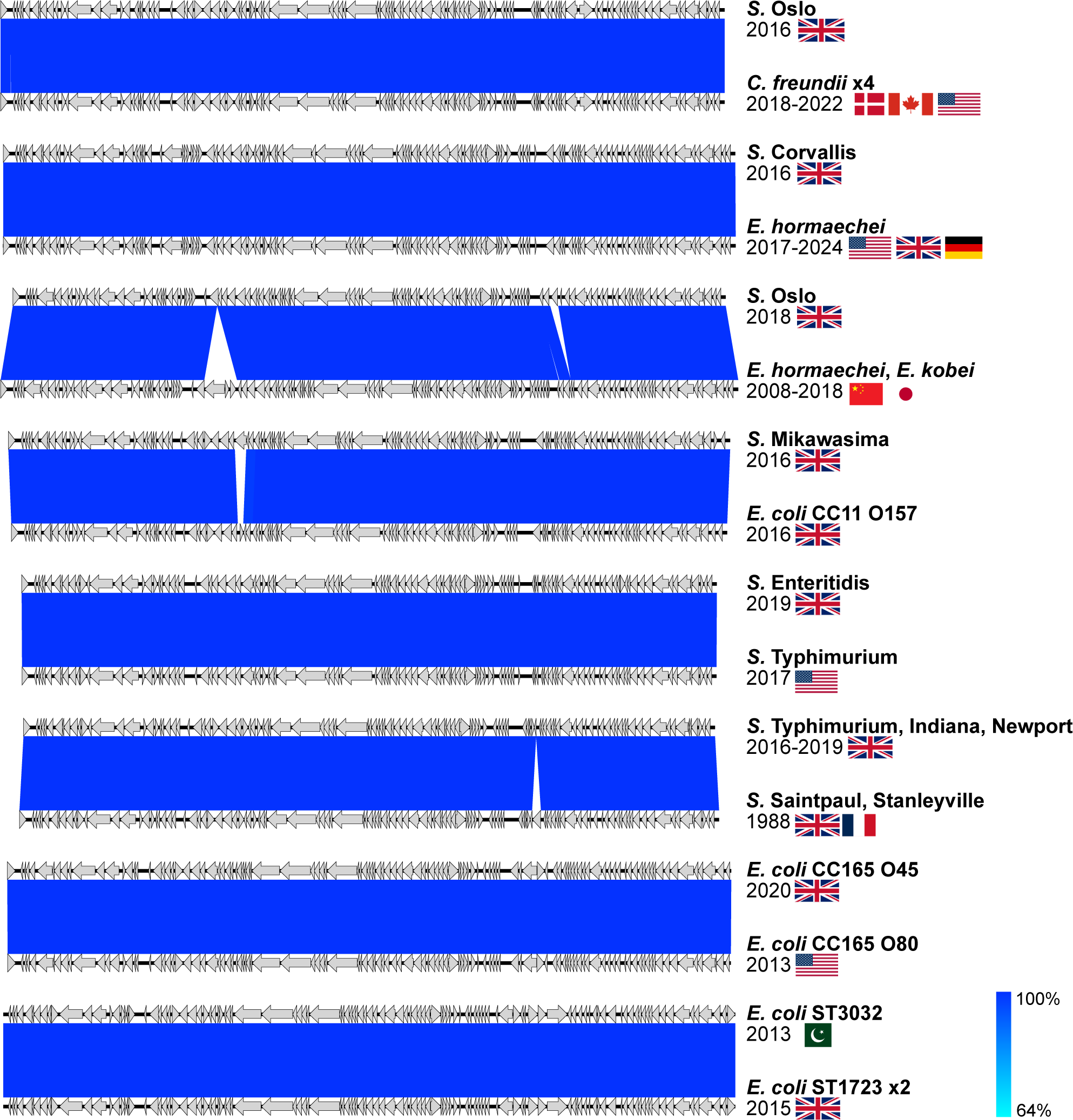
SSU5 group phage-plasmids (P-Ps) from various species, serovars and clonal complexes that show high sequence similarity to public sequences, or with the only difference being gain/loss of insertion sequence elements.

**Fig. S10.**
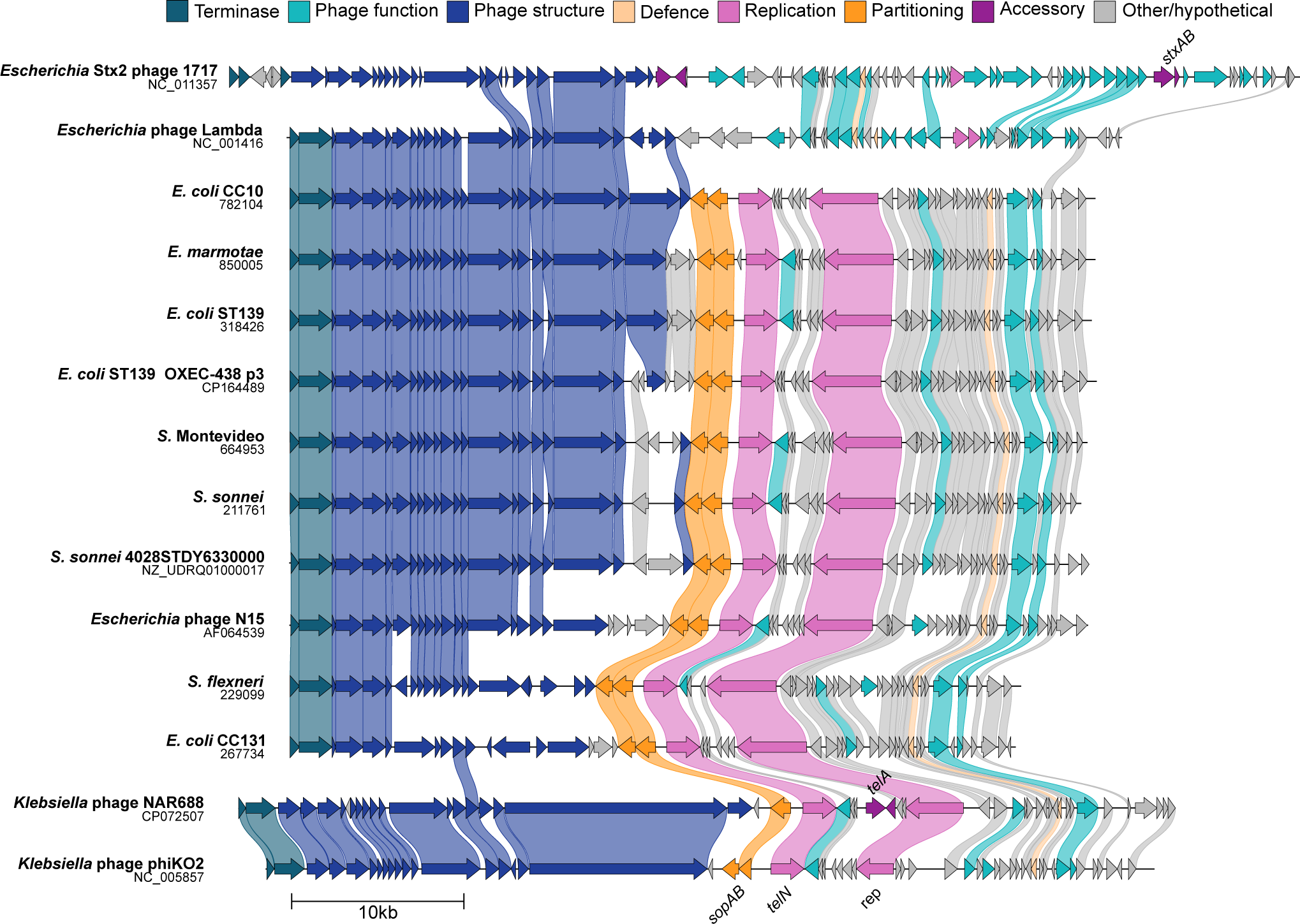
Comparison of the seven extracted N15-like sequences and public genomes showing similarities in size and layout, with genes coloured by function and linked where protein sequence identity is >30%.

